# Optimized feature gains explain and predict successes and failures of human selective listening

**DOI:** 10.1101/2025.05.28.656682

**Authors:** Ian M. Griffith, R. Preston Hess, Josh H. McDermott

## Abstract

Attention facilitates communication by enabling selective listening to sound sources of interest. However, little is known about why attentional selection succeeds in some conditions but fails in others. While neurophysiology implicates multiplicative feature gains in selective attention, it is unclear whether such gains can explain real-world attention-driven behavior. To investigate these issues, we optimized an artificial neural network with stimulus-computable, feature-based gains to recognize a cued talker’s speech from binaural audio in “cocktail party” scenarios. Though not trained to mimic humans, the model matched human performance across diverse real-world conditions, exhibiting selection based both on voice qualities and spatial location. It also predicted novel attentional effects that we confirmed in human experiments, and exhibited signatures of “late selection” like those seen in human auditory cortex. The results suggest that human-like attentional strategies naturally arise from optimization of feature gains for selective listening, offering a normative account of the mechanisms—and limitations—of auditory attention.

## Introduction

Organisms often base behavior on one of many objects in their environment. This ability typically involves endogenous (“top-down”) attention – a change in internal state that allows an organism to willfully “select” an object of interest for further processing, memory, or a behavioral response. Selective attention has been a central focus of cognitive science and neuroscience since the 1950s^1^, and much is known about both attentional abilities and their neural correlates^2-8^.

Neurophysiological observations suggest a mechanistic account of attentional selection as multiplicative gains that enhance the perceptual features of an attended object. Specifically, attention to particular features tends to cause the responses of neurons tuned to those features to be scaled upwards, enhancing the representation of objects containing those features^9-12^. Other types of effects on neural tuning^12,13^ can be explained by combining multiplicative gains with normalization^14^. Consistent with such findings, human neuroscience studies show attentional enhancement of attended sound sources^15-20^. However, it remains unclear whether feature-based multiplicative gains are sufficient to account for real-world attention-based abilities, in part because we have lacked working models of attention that can be evaluated in everyday settings. In particular, although computational models of sensory systems have made notable advances in being able to account for some types of human judgments of images and sounds^21,22^, they have thus far largely not incorporated attentional mechanisms (see Discussion).

Speech comprehension is one setting in which selective attention is essential in daily life. We routinely must understand what someone is saying despite the presence of other concurrent talkers (the “cocktail party problem”)^1,23,24^. Human attentional selection of one voice among others has been demonstrated in many settings^25-28^ and is known to depend on the features of individual sound sources – both spatial locations and voice qualities such as pitch. However, human attentional selection can be prone to failure in some settings, as when a distractor voice shares features with the target voice^26,29^. The reasons for these failures remain poorly understood. For instance, it has been unclear whether human selection errors represent suboptimal strategies, or whether such errors are largely inevitable given the feature overlap between human voices.

We used the cocktail party problem^23,24^ as a setting in which to explore computational accounts of feature-based attention. We tested whether a task-optimized model equipped with multiplicative gains applied to sensory representations would replicate human selective listening behavior. We assumed gains to be functions of a memory of the attentional target’s properties, and used a task in which the target properties could be estimated from prior exposure to a target talker’s voice. We found that models with such multiplicative gains that were optimized to recognize the words of a cued talker reproduced a wide range of characteristics of human auditory attention, including its sensitivity to both spatial and nonspatial aspects of sound, and occasional failures of selection. They also predicted two previously undocumented traits of human attentional selection that we subsequently confirmed experimentally.

The results indicate that multiplicative gains, when combined with standard filtering, pooling, and normalization operations, are sufficient to account for both the successes and failures of human feature-based attention. The framework we propose is general, and should be applicable to any behavior involving feature-based attention.

## Results

### Feature-based attention task

To study feature-based attention, we used a task in which a listener first heard an excerpt of a target voice (the “cue”), then heard a mixture of the target voice with a distractor voice, and then reported the middle word spoken by the target voice (Fig. 1a). The excerpt of the target voice used for the mixture was different from that used for the cue, such that the cue indicated the target talker’s voice and spatial location, but not the words spoken by the target talker in the mixture. To solve the task, the listener had to use properties of the cue (its voice properties and/or spatial position) to select the target voice from the mixture. The task captured key aspects of attentional selection in the wild by requiring selection based on the memory of a sound source, but its stereotyped form enabled training and behavioral benchmarking at a large scale. Depending on the experiment, sounds could be presented over headphones, with the same audio signal to both ears (eliminating any spatial cues), or from different spatial locations (with each ear receiving a different audio signal).

**Figure 1.**
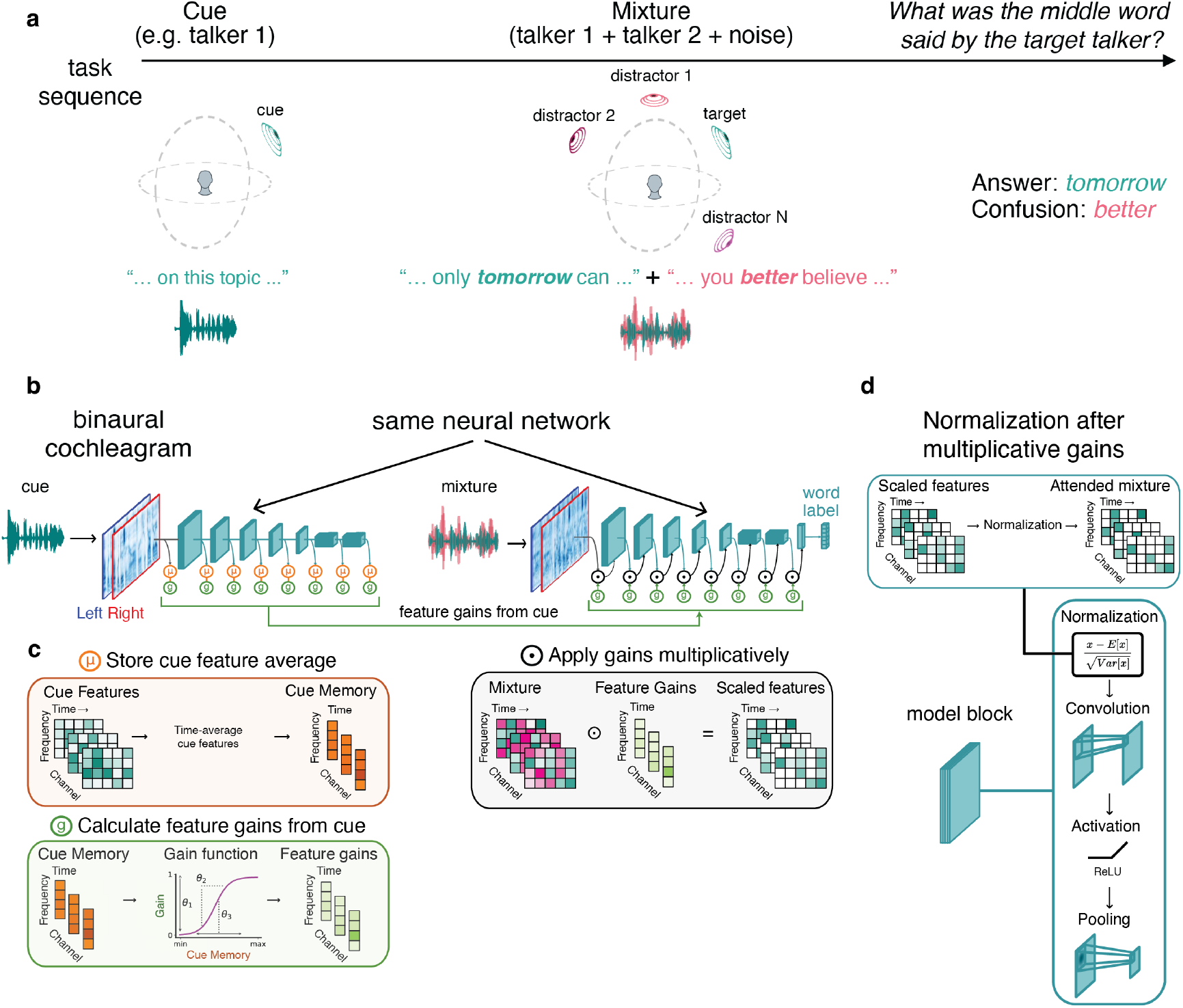
Task and model architecture used to study attention. **a**. Selective listening task. Listeners and models first heard an excerpt of a target talker’s voice (the “cue”), then heard a mixture of a different excerpt of the target talker’s speech superimposed with other sounds (the “mixture”), and then reported the word uttered by the target talker at the mid-point of the excerpt. **b**. Computational framework for modeling attention. A feedforward model of the auditory system (blue), comprised of a simulation of the cochlea followed by a neural network, took binaural audio as input and produced word labels as output. Each convolutional block of the model consisted of layer normalization, convolution, rectified linear activation, and pooling operations. When processing the mixture, the activations of the auditory model were multiplied by scalar gains (green). The gains were determined by sigmoidal functions that operated on the average activations of a cue stimulus (orange), obtained by passing the cue stimulus through the same model of the auditory system. **c**. Example application of multiplicative gains in model. Intuitively, the sigmoidal gain functions should enable gains to be high for features that have high activations in the cue, allowing these features to be passed through the auditory system, enhancing the representation of the target talker. **d**. Operations within a convolutional block of the model. Feature gains were applied immediately prior to the beginning of the subsequent model block. Each block began with a normalization operation, such that multiplicative gains were followed by normalization.

### Model optimization

We used supervised deep learning to build a model that could perform this task. Models were trained to report the middle word in a 2-second speech excerpt spoken by a cued talker within a mixture of talkers. Both cue and mixture were presented as binaural audio. During training, audio was spatially rendered at locations within simulated reverberant rooms using head-related transfer functions to reproduce the spatial cues available to humans. To emulate the variety of real-world scenarios encountered by humans, each mixture in the training data varied in the number of distractor sources, whether distractors were speech or non-speech, the relative intensities of each source, their spatial configuration, and the dimensions of the room they were heard in. The cue was always an excerpt of the target voice in isolation, presented at the same location as the target in the corresponding mixture. Models classified the middle word within the target excerpt (out of 800 possible word classes).

### Model architecture

Our main model architectures (referred to as “feature-gain models”) incorporated learnable attentional gain functions into a deep neural network model of the auditory system (Fig. 1b). The neural network took simulated cochlear representations of an audio waveform as input. Feature-based attention was implemented with parameterized sigmoidal functions (green box in Fig. 1c) which mapped the model’s internal representation of the cue to scalar multiplicative gains. These gains were then applied multiplicatively to the mixture representation.

The model architecture instantiated the hypothesis that attention is mediated by multiplicative gains that are deterministic functions of the features in a mental representation of the target of attention. To compute attentional gains, the cue stimulus for a trial was first passed through the auditory system model (blue model stages in Fig. 1b). The activations of each model stage were then time-averaged and stored to obtain stage-dependent memory representations of the cue (orange stages in Fig. 1b&c). Gains were computed as a sigmoidal function of the corresponding memory representations (features with high activations in the cue yield high gains), which were then multiplied with the activations of the mixture representation during the forward cascade of the model (Fig. 1b). Intuitively, features that are present in the cue should result in high gains, passing those features through the model when they occur in the mixture. The sigmoidal functions should enable models to learn how strongly to modulate the features at each model stage to select the target from the mixture. We used a single set of shared sigmoid parameters per model stage (we found this was sufficient to yield good performance). Gain function parameters and model features were jointly optimized, such that models should learn to prioritize information that enables selection of the target voice content.

We note that the application of feature gains happened immediately prior to each convolutional “block”, which always began with a normalization operation. The model thus instantiated the key components of the normalization model of single-unit attentional effects^14^, but embedded in a hierarchical system of learnable gains and features.

To ensure the results generalized across the details of the model architecture, we trained 10 different feature-gain models with different architectures (Supplementary Table 1). Results for the feature-gain models are the average over these 10 architectures. In a later section we test the extent to which the feature-gain architectural assumption was important to the results.

### Human-model behavioral comparisons to assess attentional strategies

Because the models receive binaural audio signals, they should be able to learn to use both spatial cues and voice timbre cues if they are useful in solving the task. However, because the models were optimized only for word recognition, they were not explicitly constrained to reproduce human-like strategies, and were free to learn any solution that yielded good performance. To assess whether the model reproduced the properties of human attention, we conducted a set of experiments to characterize the model’s behavior and to compare it to that of humans.

### Model replicates human cocktail party performance in monaural conditions

We first ran human participants in a behavioral benchmark (Experiment 1) containing a set of diotic listening conditions (i.e., in which the same audio waveform was presented to the left and right ears, eliminating binaural spatial cues). We then simulated the same experiment on the model. Evaluation stimuli were new to the model (and to participants). As shown in Fig. 2, the model approximately replicated both the overall performance of human listeners and the dependence on signal-to-noise ratio and distractor type (see Supplementary Fig. 1 for results for the 10 individual model architectures, and Supplementary Fig. 2 for plots of data distributions).

**Figure 2.**
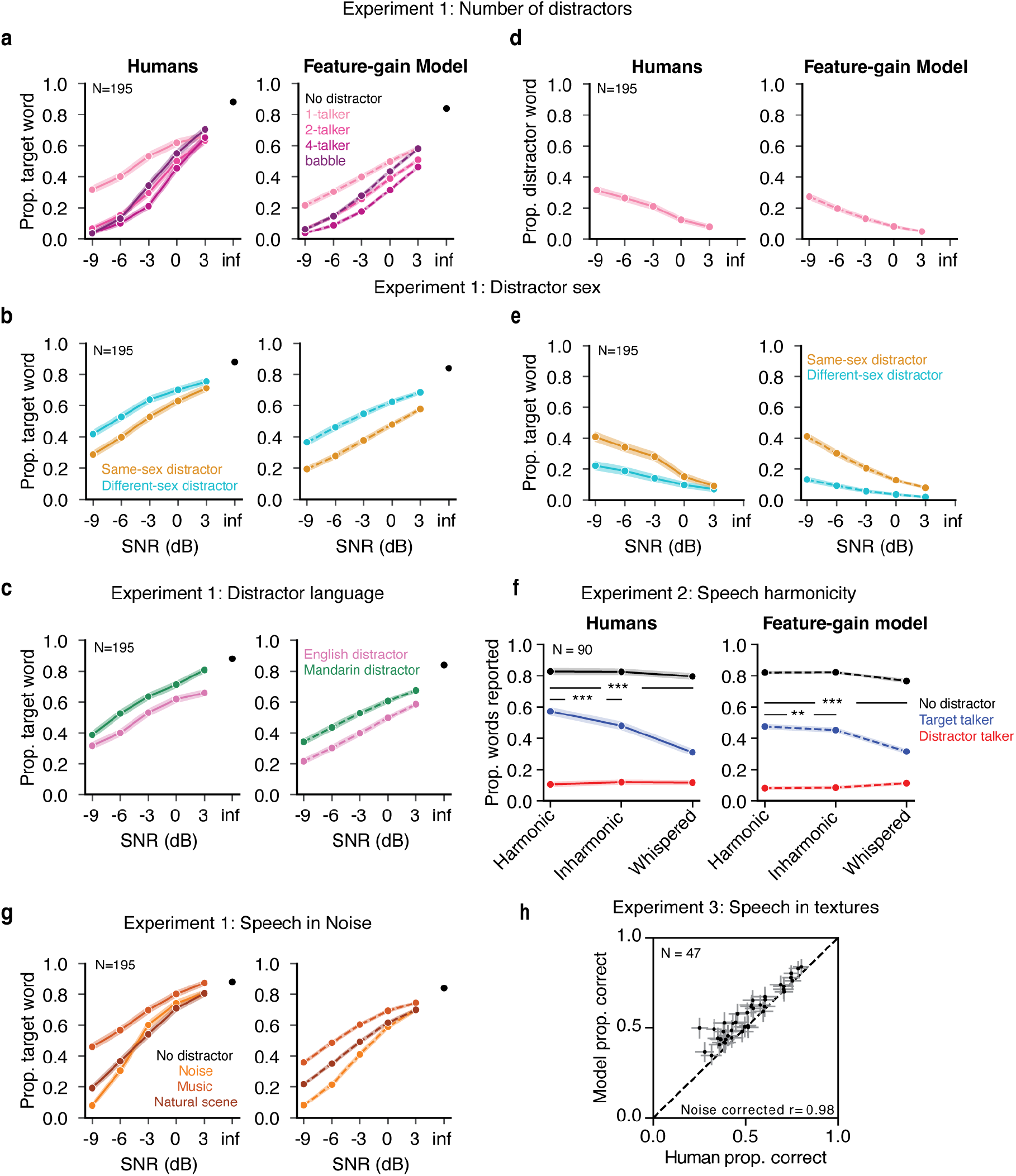
Comparison of human and model attentional selections in monaural conditions. **a**. Human and model performance vs. signal-to-noise ratio for speech distractor stimuli (Experiment 1). Here and in other panels, error bars plot SEM. Note that there is only one data point for the infinite signal-to-noise ratio condition, because the distractor is zeroed out in this condition, making the different distractor conditions equivalent. **b**. Effect of same-vs. different-sex distractor talkers (Experiment 1). **c**. Effect of foreign language distractor talkers (Experiment 1). **d**. Human and model confusions, plotted vs. signal-to-noise ratio (Experiment 1). Confusions occurred when the participant erroneously reported the word uttered by the distractor talker. **e**. Human and model confusions plotted separately for same- and different-sex distractor talkers (Experiment 1). **f**. Effect of talker harmonicity (Experiment 2). Note that the rate of confusions (reports of words uttered by the distractor talker) are low because the SNR was always 0 dB, at which there are few confusions in Experiment 1 as well. **g**. Effect of non-speech distractor sounds (Experiment 1). **h**. Human and model performance across a large set of distractor noises (Experiment 3).

Previous models of word recognition have replicated patterns of intelligibility across different types of noise^22,30^. However, because they lacked the ability to use a cue to selectively report one of several talkers, they were unable to perform at human levels in the presence of a single distractor talker, because without a cue, the task is ill-defined. As a result, it was not clear a priori that a model would be able to match human behavior in this regime. However, the feature-gain model performed similarly to humans on trials including only the target and a single distractor talker (1-distractor condition in Fig. 2a). The similarity to human performance in this condition indicates that the model succeeds at attentional selection on par with human listeners. Human and model performance was robust to variation in cue duration (Supplementary Figure 3), demonstrating some generality in the model’s behavior despite the stereotyped structure of the training examples.

### Model replicates effects of distractor language and sex

To assess whether the model was sensitive to some of the same aspects of speech that influence human attentional selection, we reanalyzed the one-distractor condition of Experiment 1. Human attentional selection is known to improve when target and distractor talkers differ in sex, in part because of the acoustic differences this entails^26^. Selection is also better when distractor talkers speak a language unfamiliar to the listener^31,32^. Native English-speaking human participants in Experiment 1 replicated both of these effects, showing higher word recognition performance with different-sex distractors (Fig 2b) and Mandarin language (Fig 2c). The model also exhibited both effects, indicating the model learned to rely on some of the same cues as humans.

### Model exhibits human-like failures of attention

Human attention is also known to demonstrate systematic failures. Even when attempting to attend to a target talker, humans sometimes mistakenly report what was said by a distractor talker instead. The root causes of these failures are not clear.

To quantify selection failures, we re-analyzed the one-distractor condition of Experiment 1, measuring how often listeners reported words in the distractor utterance (“confusions” of the target and distractor). The overall confusion rate was low, but increased at lower SNRs (*f*(4,776) = 51.707, *P* < 0.0001, 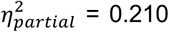), and when the target and distractor were the same sex (*f*(1,194) = 89.108, *P* < 0.0001, 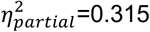; Fig. 2d&e). The model exhibited quantitatively similar effects (Fig. 2d&e). This result suggests that some selection failures are an inevitable consequence of target-distractor feature similarity. We note that humans made slightly higher rates of confusions than the model, which could potentially also reflect lapses or failures of executive function, which undoubtedly contribute to human errors in some settings.

### Model replicates effects of speech harmonicity

Human attentional selection has also been shown to depend on whether the constituent frequency components of speech signals are harmonically related^33^. We ran an additional experiment to measure this effect using our task (Experiment 2), and simulated the same experiment on the model. Target, distractor, and cue signals were resynthesized to be harmonic, inharmonic, or whispered (always of the same type, e.g. all harmonic, all inharmonic, or all whispered). Without a concurrent distractor talker, human word recognition was similar for resynthesized harmonic, inharmonic, and whispered speech (Fig. 2f; see Supplementary Fig. 4 for plots of data distributions). But with a concurrent distractor, human recognition was modestly impaired for inharmonic speech, and substantially impaired for whispered speech, producing an interaction between the effect of speech harmonicity and single vs. mixtures of talkers (F(2,178) = 51.021, p < 0.0001, 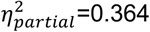), as in prior work (Fig. 2f). The model qualitatively reproduced these effects, also showing an interaction between speech harmonicity and single vs. mixtures of talkers (F(2, 18) = 116.151, p < 0.0001, 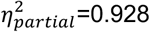), driven by worse performance for inharmonic (*t*(9) = 5.752, *P* = 0.0003, Cohen’ s d = 1.819), and whispered (*t*(9) = 15.142, *P* < 0.0001, Cohen’ s d = 4.788) speech compared to harmonic speech in concurrent distractor conditions.

### Model replicates human speech-in-noise performance

The model also reproduced human-like patterns of performance across different types of noise distractors, like previous models^22,30^. In Experiment 1, both humans and the model demonstrated better performance with noise distractors compared to speech distractors, and for some types of noise compared to others (Fig. 2g). Fig. 2h shows an additional experiment (Experiment 3) comparing the model and human listeners across a large set of different types of (non-speech) noise^22^, further demonstrating the model’s human-like dependence on noise type.

### Models replicate signatures of human spatial attention

Humans also benefit from spatial separation between sources, an effect often termed “spatial release from masking”. This benefit is thought to be partly attentional in origin, as it depends in part on knowing where to listen^27^. Because our model was trained on binaural audio rendered from sounds at different locations, it could in principle learn to use spatial information to aid selection. However, because models were only optimized to recognize words, it was not obvious if spatial attention would emerge as a listening strategy.

To test if the models exhibited human-like effects of spatial separation, we evaluated performance as a function of target-distractor separation in azimuth. We replicated a previously published experiment in human listeners^34^ in which performance was measured with distractors placed symmetrically in azimuth (Fig 3a). This design has the advantage of preserving the signal-to-noise ratio (SNR) at both ears as the distractors are moved away from the target talker, such that an advantage of spatial separation cannot be explained merely by increased SNR at one of the ears. As in the human experiment, we summarized performance at each spatial separation with a threshold (the signal-to-noise ratio granting 50% of ceiling performance). The model displayed thresholds that varied with spatial separation on par with humans (main effect of offset for model, F(4,36) = 426.57, p<0.0001, 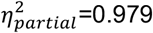; Fig. 3b), suggesting that it also made use of spatial location for selection.

**Figure 3.**
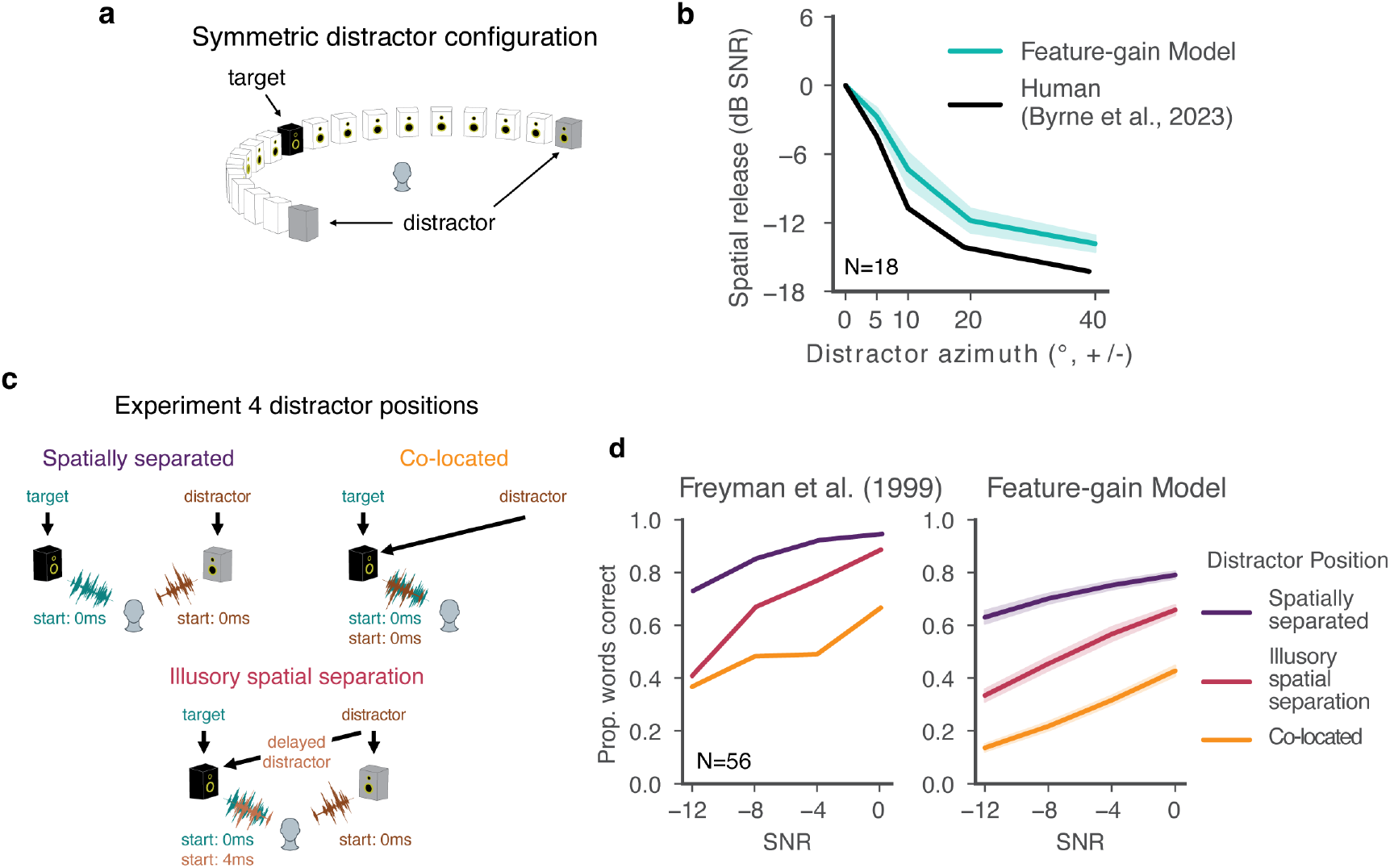
Comparison of human and model spatial attention. **a**. Stimulus setup for Experiment 4. The target talker was positioned in front of the listener, with distractor talkers positioned symmetrically on either side. Word recognition performance was measured as a function of signal-to-noise ratio for different target-distractor spatial offsets. **b**. Results of Experiment 4. Graph plots spatial release from masking, measured as the decrease in speech reception thresholds from the co-located condition where distractor talkers were also positioned directly in front of the listener. Human data were scanned in from original publication^34^ and replotted. Error bars plot 2 SEM for the Feature-gain model (error bars for Humans not available in original publication). **c**. Stimulus setup for Experiment 5. Speech signals were played from two speakers. Target and distractor could be spatially separated, co-located, or given illusory spatial separation by virtue of the precedence effect. In this latter condition the target and distractor talker were both played from the left speaker, and a second copy of the distractor speaker was played from the right speaker. The co-located copy of the distractor was slightly delayed relative to the separated copy, inducing the illusion that the distractor came from the right speaker. This effect is commonly thought to reflect a localization strategy adapted to cope with environmental reflections. Reflections typically arrive from directions distinct from the true direction of the source from a listener, but are delayed due to the increased path length compared to the sound that arrives directly from the source. The human auditory system appears to suppress localization cues from delayed sound components that are likely to be due to reflections, presumably to achieve more robust localization in real-world conditions. **d**. Results of Experiment 5. Graph plots word recognition vs. signal-to-noise ratio for the three different spatial configurations. Error bars plot 2 SEM for the Feature-gain model (error bars for Humans not available in original publication).

### Model replicates human advantage with illusory spatial separation

Another compelling demonstration of spatial attention in humans comes from the fact that human listeners benefit from illusory separation of sources. Such illusory separation can be mediated by the “precedence effect”^35,36^, in which a sound played in rapid succession from two locations is heard to come from the first location, likely a consequence of a strategy for robust localization in the presence of reflections^37^. Specifically, a distractor signal co-located with a target signal (Fig. 3c, top right) impairs speech recognition relative to when it is spatially separated (Fig. 3c, top left). But if a second, spatially displaced copy of the distractor signal is played along with the co-located signal, human performance improves provided the displaced copy begins shortly before the co-located copy. The presumptive explanation is that the co-located copy is interpreted by the brain as a reflection, such that the distractor is represented as coming from the displaced location, and can be filtered out with spatial attention. To test whether the model similarly benefitted from illusory spatial separation, we simulated a previously published experiment^38^ contrasting the benefit of illusory and true separation (Fig. 3c). As shown in Fig. 3d, model performance benefited from illusory separation in a qualitatively similar way as in humans (main effect of illusory separation vs. co-located, F(1,9) = 1224.917, p<0.0001, 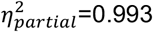). As in models optimized for sound localization in realistic environments^37^, the model presumably learns representations of sound location that are adapted to the presence of reflections (potentially by suppressing the localization cues of late-arriving sound akin to the biological auditory system^39^), allowing it to benefit from the illusory spatial separation in this experimental setting.

### Model predictions of human behavior

There are many demonstrations of auditory spatial attention in humans, but they have mostly been restricted to a fairly consistent set of spatial configurations. One benefit of a stimulus-computable model is that experiments on the model are inexpensive, such that the model can be used to screen large numbers of experimental conditions to search for interesting effects. Such effects can then be tested in humans, serving both to further characterize human perception, and as strong tests of the model. With these goals in mind, we ran the model on all possible combinations of target-distractor locations, summarizing the effect on performance in Fig. 4a (see Supplementary Fig. 5 and 6 for full set of results). Two effects stood out from this exhaustive screen, which we subsequently probed for in humans.

**Figure 4.**
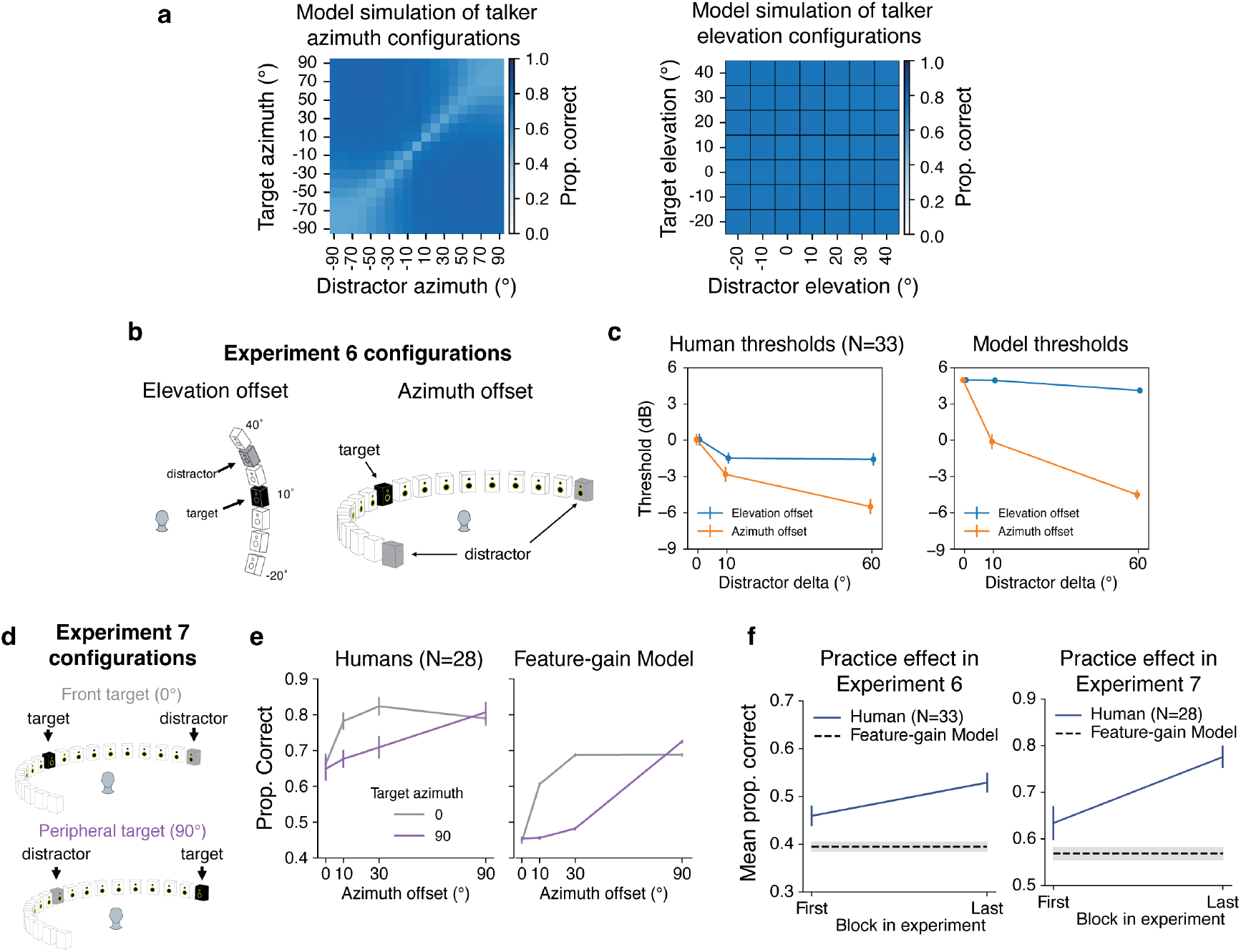
Model predictions of human spatial selection. **a**. Model word recognition performance across all possible combinations of target and distractor locations. Results are averaged over elevation and the front and back hemifields (left) or azimuth (right) for ease of visualization. Model results suggest larger benefits from azimuthal separation than elevation separation (note that there was almost no effect of separation in elevation, such that the right panel is uniform), and a “spotlight” that varies in width, being narrow for targets at the midline and wide for targets at peripheral locations. **b**. Spatial configurations tested in Experiment 6. To eliminate the possibility that an advantage from azimuth offset might be due to changes in signal-to-noise ratio in the input to an ear, the azimuth offset condition presented two distractor talkers, symmetrically located about the target talker position. Elevation offset condition presented two distractor talkers at one elevation (constrained by the locations that were possible given the speaker array). **c**. Results of Experiment 6. Humans and models show larger benefit of spatial separation in azimuth than in elevation. Here and in panels e and f, error bars plot SEM. **d**. Spatial configurations tested in Experiment 7. Target talkers were positioned at either 0 or 90 degrees, with distractors at one of four offsets. **e**. Results of Experiment 7. Error bars plot SEM. **f**. Practice effect evident in human performance in Experiments 6 and 7. Graphs plot human performance in the first and last block of each experiment, compared to overall model performance.

### Model predicts horizontal/vertical asymmetry in spatial selection

When we exhaustively tested target-distractor configurations on the model, target word recognition increased as a function of the distractor spatial offset, as expected. However, offsets in the vertical direction produced much less masking release than the same offsets in the horizontal direction (Fig. 4a). Such differences are plausibly due to the different cues involved in the two types of offset – horizontal offsets are mostly signaled by differences between the ears, whereas vertical offsets are mostly signaled by monaural cues. It seemed plausible that monaural cues might be less robust to the presence of concurrent sources (see Supplementary Fig. 7). This issue has been addressed in a handful of previous studies^40-45^, but with mixed results, and to our knowledge, the effect of horizontal and vertical offsets had never been compared in the same setting.

To test for analogous effects in humans, we ran an experiment with human participants measuring speech reception thresholds (i.e., the signal-to-noise ratio permitting average performance of 50% correct) for different target-distractor offsets in azimuth and elevation (Fig. 4b; Experiment 6; see Supplementary Fig. 8 for plots of data distributions). As expected, thresholds decreased as azimuthal distractor offsets increased, replicating known effects of spatial release from masking. However, the benefit was much weaker in elevation (Fig. 4c; significant interaction between direction of offset and extent of offset, p < 0.0001, using a non-parametric test for interaction, see Methods), confirming the model prediction.

### Model predicts central/peripheral differences in width of spatial “spotlight”

A second effect evident in Fig. 4a is that the spatial separation between target and distractor needed to obtain a benefit is much less for targets at the midline (0°) than for targets in the periphery (±90°). See Supplementary Fig. 9 for further analysis of this effect for both speech and noise distractors. This effect is plausibly due to the higher acuity of localization at the midline^46-48^, which in turn is thought to relate to the derivative of binaural cues with spatial position (see Supplementary Fig. 7). However, it is not obvious that spatial acuity for isolated sources should directly translate to the “width” of spatial attention effects. To our knowledge, this issue had not been previously examined in humans.

We ran an experiment with human participants measuring the effect of masking release as a function of distractor offset, for both centrally and peripherally positioned targets (Fig. 4d; Experiment 7). The model was run on a simulation of the same experiment. As shown in Fig. 4e, the qualitative difference between central and peripheral targets seen in models was also evident in humans: the distractor had to be spatially offset by a much larger amount for peripheral targets to yield the same selection benefit seen at smaller offsets centrally positioned targets, producing a significant interaction between the effect of target-distractor offset and target position (F(3,81) = 6.731, p=0.0004, 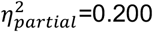).

We note that in both Experiments 6 and 7, human performance was overall somewhat higher than model performance. An analysis of human performance over the course of both experiments showed evidence of a practice effect in humans. Specifically, human performance was similar to model performance at the start of the experiment, but got better over time (Fig. 4f). A plausible explanation is that human participants adapted to the room in which the experiment was conducted^49,50^, boosting their performance (the model has no such ability as its weights were frozen following training, and did not change over the course of simulated experiments).

### Models exhibit signatures of late selection

A signature of human auditory attention to speech is the enhancement of the neural representation of a target source at relatively late stages of the presumptive auditory hierarchy, such as non-primary auditory cortex^18,19,51-54^. However, it remains unclear why attentional enhancement might be limited to particular processing stages. In principle, the stage at which enhancement occurs could be determined by anatomical constraints on top-down connections. But they might also simply reflect the stage at which features are most useful for selection. Task-optimized models provide one way to gain insight into this issue, because they reveal an optimized solution that can be compared to that of the brain.

To assess the locus of attentional enhancement, at each model stage we measured correlations between the activations of target-distractor mixtures and the activations of either the target or distractor alone^16^ (Fig. 5a). Attentional enhancement of the target should result in target-mixture correlations being higher than distractor-mixture correlations. Differences between target-mixture and distractor-mixture correlations were largest for configurations with spatially separated target and distractor signals, but in all cases became pronounced only at later model stages (Fig. 5b), indicating that enhancement occurs relatively late in the model. See Supplementary Fig. 10 for results plotted separately for individual model architectures. The same analysis performed on a model with randomly initialized weights did not show evidence for stage-specific enhancement (Fig. 5c), indicating that this result is not an inevitable consequence of the model architecture. This result is qualitatively consistent with human neuroscience evidence^18,19,52-54^, and suggests that the solution arrived at in the brain could reflect the representational locus at which enhancement is effective in enabling speech recognition.

**Figure 5.**
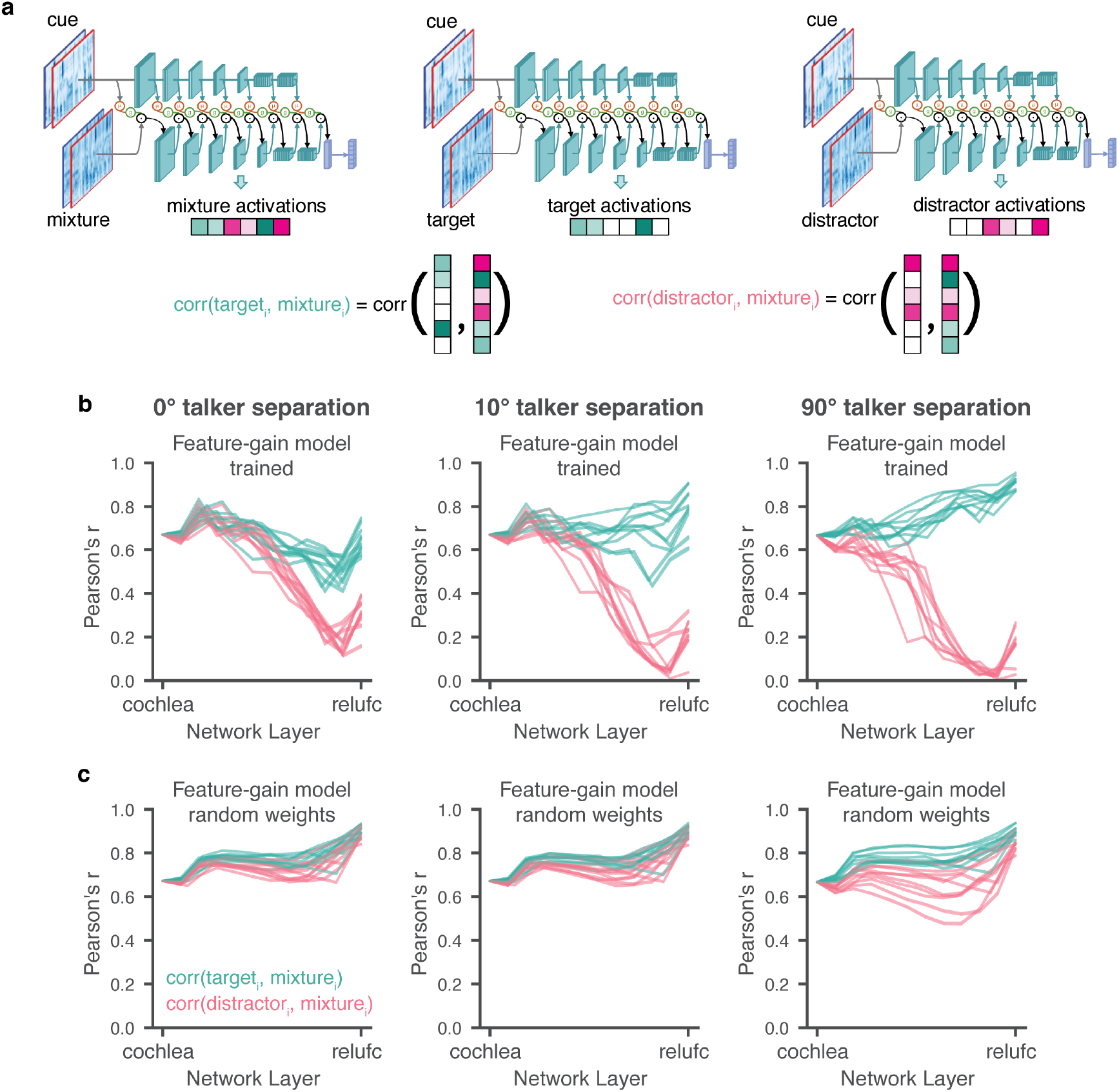
Stage of attentional selection. **a**. Explanation of stage-of-selection analysis. Target, distractor, and mixture excerpts were passed through the auditory model, with feature-gains derived from a cue matching the target. At each model stage, the activations of the target and distractor were correlated with the activations of the mixture. **b**. Task-optimized feature-gain models show signatures of target selection in late stages. Each graph plots results for a different spatial configuration of target and distractor. Each line plots results for one of 10 model architectures. In all architectures and configurations, the late-stage representation of the mixture is more similar to that of the target than that of the distractor. In b and c, error bars on results for individual models are omitted for clarity. **c**. Same as b, but for models with random weights. Note that the modest difference between the target- and distractor-mixture correlations with 90 degrees separation likely reflects the large different in SNR that occurs in this condition within an ear, such that random features that by chance have greater input from one ear than the other are sufficient to achieve a modest degree of selection.

To examine how selection relates to feature tuning, we measured model activations for a set of speech excerpts rendered at different spatial locations, and analyzed the proportion of units at each model stage that were selective for location, pitch, talker, and the middle word of the excerpt (Supplementary Fig. 11). This analysis revealed that tuning to location and talker – the two dimensions used for selection in the task – was prevalent in early model stages, with tuning for words being weak in early stages and stronger later. Overall, this tuning is consistent with the results of Fig. 5, in that tuning to the dimensions used for selection is present prior to the stages where selection is evident, such that feature gains could be used there to help select the target.

### Models lacking architecturally constrained feature gains are less human-like

The inclusion of gain functions in the model instantiates an inductive bias on how attention could work. To investigate the importance of this inductive bias, we trained several alternate versions of the model (Fig. 6a). A “baseline” model removed the constraint of explicit gain functions entirely, instead receiving the mixture and cue as separate input channels, with model weights again optimized for recognition of the target word in the mixture. To test the importance of the application of gains followed by normalization, we also trained a feature-gain model in which this ordering was reversed, with gains applied after normalization (“gain-after-normalization”). We also ran alternate architectures in which attention was constrained to operate only on particular stages. An “early-only” model had gains applied only at the cochleagram stage, whereas a “late-only” model had gains applied only at the final fully connected model stage. Because each model stage had a single shared sigmoid gain function defined by only three parameters, the early-only, late-only, and feature-gain models had nearly the same number of parameters (differing by 21 parameters out of 62.6 million). The baseline model had slightly more parameters (just over 69 million) due to the extra input channels, but the models were overall similar in complexity.

**Figure 6.**
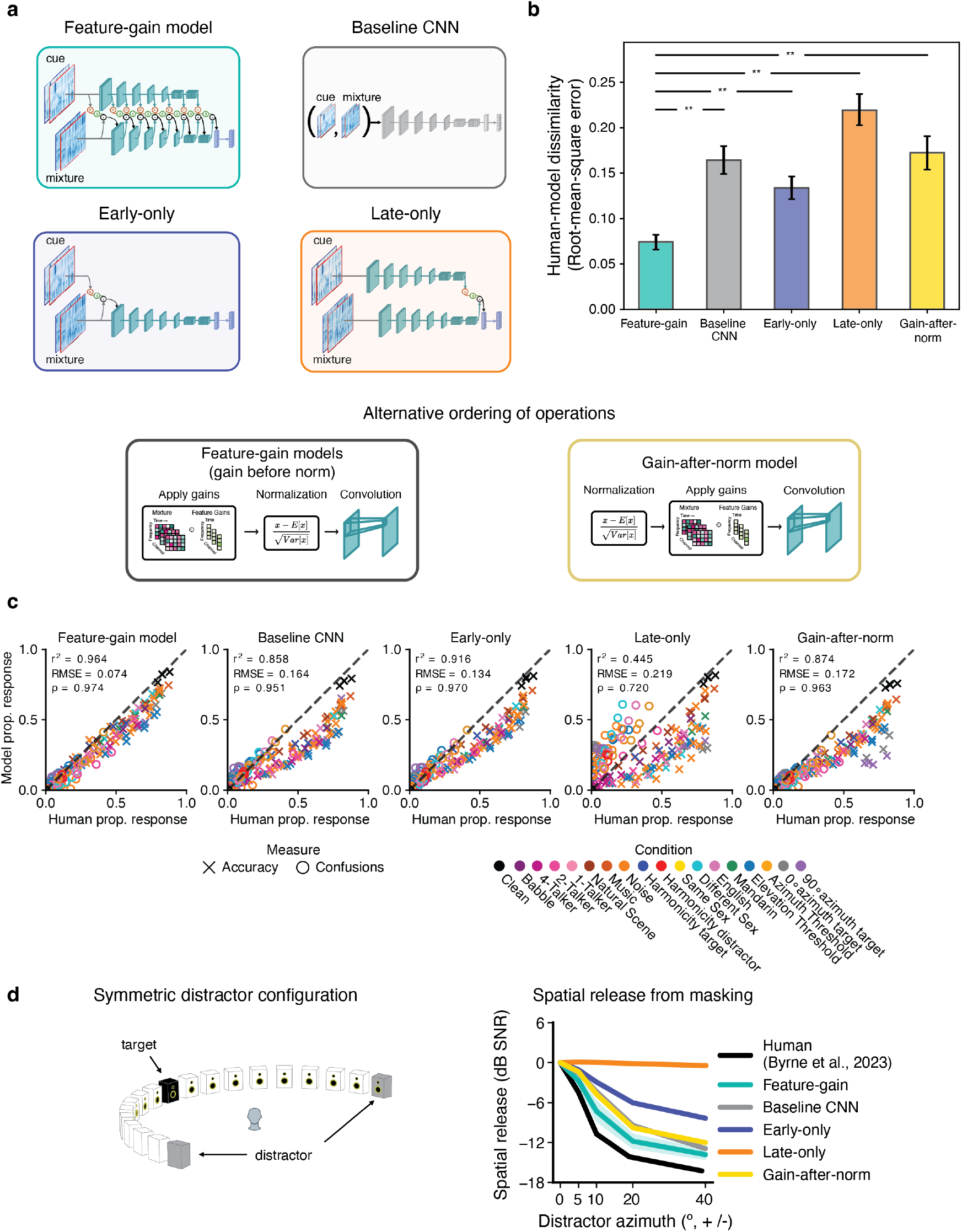
Dependence of attentional selection on model architecture. **a**. Alternative model architectures: i) the main feature-gain architecture; ii) a baseline architecture in which the cue was supplied to the model as an additional input channel, without explicit feature gains; iii) gain-after-normalization architecture; iv) early-only architecture in which feature gains were applied only at the cochleagram; v) late-only model in which feature gains were applied only at the last convolutional stage. **b**. Aggregate measure of human-model behavioral dissimilarity for each model architecture. Graph plots root-mean-squared error between human and model performance in all experimental conditions. Error bars plot 95^th^ confidence intervals obtained via bootstrap of human-model similarity. **c**. Scatter plots of model vs. human performance in each of the experimental conditions from Figures 2-4. Each of the alternative architectures produces a worse match to human performance in some conditions. **d**. Performance of alternative model architectures (feature-gain, baseline, early-only, and late-only models) as a function of target-distractor separation in azimuth for symmetrically positioned distractors (Experiment 4). Models are compared to human data from Byrne et al. (2023)^34^. The y-axis plots spatial release from masking (the decrease in speech reception thresholds compared to the co-located distractor condition) at each tested azimuthal offset. Each of the alternative models displayed less human-like spatial release from masking than the feature-gain model. The late-only model showed no spatial masking release (orange), and the others exhibited degraded masking release compared to the feature-gain model.

After optimizing each of these alternative models we ran them on each of the previously described experiments, measuring the proportion of target words correctly reported in each condition as well as the proportion of confusions in single-distractor conditions. We compared each model’s pattern of performance across conditions to that of humans, using both root-mean-squared error and correlation metrics. These two metrics are complementary: RMS error metric captures differences in both absolute and relative performance, whereas the correlation metric isolates differences in relative performance. A good model should match both absolute performance and relative performance differences between conditions, and should match humans as well or better than other models on all metrics. We also used a mutual information metric (Supplementary Fig. 12), which yielded similar conclusions. As an overall summary measure, we jointly analyzed both the proportion correct in each condition, and the proportion of confusions in each single-talker-distractor condition.

Overall, the feature-gain model explained much of the variance in human performance across all experiments, and the explanatory power depended on the architectural constraints provided by the feature gains (Fig. 6b; see Supplementary Fig. 13 for results for each individual model architecture). Each alternate architecture showed significantly lower human-model similarity (root-mean-squared error: feature-gain vs baseline p=0.002, difference of means = −0.070; feature-gain vs gain-after-normalization p=0.002, difference of means = −0.078; feature-gain vs early-only p=0.002, difference of means = −0.040; feature-gain vs late-only p=0.002, difference of means = −0.125; Pearson’s *r*^2^: feature-gain vs baseline p=0.002, difference of means = 0.093; feature-gain vs gain-after-normalization p=0.002, difference of means = 0.007; feature-gain vs early-only p=0.002, difference of means = 0.036; feature-gain vs late-only p=0.002, difference of means = 0.507). Inspection of the pattern of proportion correct reveals that the alternative models tended to perform worse than the feature-gain model, but were also less correlated with humans across conditions. In particular, there was weaker spatial release from masking for each alternative model. See Supplementary Fig. 14 for explicit comparison of model performance.

These results confirm that the architectural bias of multiplicative gains helps to reproduce human-like attentional behavior. We emphasize that the results do not preclude the (likely) possibility that a sufficiently large baseline model, trained on enough data, could reproduce the performance of the feature-gain model (potentially by learning to replicate the effect of the gains). We view the feature gains as analogous to other architectural motifs (e.g. convolution) which in principle could also be learned by a less structured model. We also emphasize that the alternative models were fully optimized with the alternative architecture, and were free to learn an entirely different feature hierarchy that allowed them to maximize their performance given the constraint of gains in particular locations (or no explicit gains at all). It is thus perhaps not surprising that these models were able to perform well above chance. The success of the architectural constraint in replicating human behavior nonetheless provides support for its role in biological attention.

## Discussion

We investigated feature-based attention using stimulus-computable task-optimized models of the cocktail party problem. We augmented standard feedforward neural network architectures with memory-driven multiplicative feature gains, and optimized the models to report the word spoken by a cued talker given only binaural audio. The resulting models replicated the phenotype of human auditory attention, correctly reporting a cued talker’s speech at comparable levels to humans, and exhibiting performance variation across conditions like that of humans. In particular, models exhibited human-like advantages for different-sex and different-language distractors, harmonic voices, and both real and illusory spatial offsets between target and distractor talkers. We used the models to sample target-distractor spatial arrangements more exhaustively than has been possible in human listeners, and saw two notable effects that we then tested in humans. Both of these model predictions were borne out in human listeners. The model also made errors in the same settings as humans, and to around the same degree. However, model performance, and human-model similarity, were dependent on the architectural motif of multiplicative gains, being worse in models without explicit gain functions, or with gains restricted to either early or late model stages alone. These results provide support for the idea that feature-based attention can be explained by multiplicative gains, and suggest that both human successes and failures of attention reflect an optimized solution to the problem of selecting a sound source via its features. Lastly, inspection of the model representations showed evidence of late selection, providing a normative perspective on effects seen neurophysiologically.

### Relation to prior work

Although the conceptual ingredients of our model have a long history in attention research, there has been little prior work incorporating them into working models of sensory systems. Previous auditory models of the cocktail party problem have largely focused on the problem of inferring the distinct sources underlying an auditory scene, without a means to direct attention to one or more of the sources^55-61^. Previous computational work on attention has tended to either model effects of attention on neural responses^14,62^ (rather than behavior), or has considered behavioral effects in small-scale models and simple tasks^63-68^. The main previous attempt to test the effect of multiplicative gains in a working model tested vision models on an object detection task in which four different images of objects were concatenated^69^. This study found that the application of gains proportional to a unit’s selectivity for a particular object category increased the likelihood that the model would report that category, as might be expected to occur behaviorally in humans. A related recent vision model used feedback connections as a way to emphasize particular object categories, again showing that this aided a model’s object detection in concatenations or superpositions of object images^70^. Our work builds on these efforts by a) optimizing attentional mechanisms for task performance, b)showing that a single computational framework can account for many of the known attentional phenomena in the domain we consider (attention to speech), and c) using a model to make predictions about human attentional selection (and then validating these predictions).

Other work has used models to explore other aspects of attention. For example, neural network models can be trained to guide simulated eye movements during visual search using priority maps computed from a target image, and reproduce aspects of human visual search behavior^71,72^. “Bottom-up” exogenous attentional cueing effects can also in some cases emerge naturally in task-optimized models^73^; such effects are widely believed to tap into distinct mechanisms from the endogenous effects we studied. We note that the word “attention” is also used to refer to a computational motif within transformer architectures popular in current machine learning^74^, but that this motif differs from biological selective attention in not being directed to a particular target object or sound source.

Our work was inspired by a large body of neuroscience experiments documenting neural correlates of attentional selection^18,19,52-54,75^ and suggesting that such correlates can be explained by multiplicative effects on neuronal responses^10,11^. However, such experimental findings leave it unclear whether the documented neurophysiological effects are sufficient to account for behavioral effects. Our results indicate that multiplicative gains applied at the appropriate stage of processing, with the right features, are sufficient to enable human-like auditory attentional behavior. The model results leave open the neurobiological implementation of the cue memory and feature gains, but are compatible with previous proposals for sources of feature-based attention in prefrontal cortex^76,77^, and of gain changes implemented via combinations of excitation and inhibition^78^.

One of the main debates surrounding attention involves the stage at which attentional selection occurs^52,54,79-82^. These debates originally concerned whether attention acted before or after “semantic” processing, but with the discovery of attentional effects within sensory systems, interest shifted to differences in attentional modulation between stages of sensory hierarchies^83^, and it is that setting in which we contrast “early” and “late” selection. Our results show how relatively late selection can emerge as an optimized solution for attentional selection of speech. This optimized solution could also depend on the system architecture or on other biological constraints we did not model. Our result thus does not provide a definitive explanation of the locus of selection found in the brain, but it illustrates a computational approach to understanding the issue. It also remains possible that the optimal stage of selection depends on the task^84^, which could be investigated by applying our framework to multiple tasks.

Our work builds on a large prior literature that has documented human performance in selective listening tasks with speech^25-29,31-33,38^. The model clarified these previously documented phenomena in several ways. First, the similarity of human and model performance (in particular, the similar extent and pattern of selection errors) suggests that human selection failures may partly be an inevitable consequence of feature overlap between target and distractor talkers (some are surely also due to lapses, distraction and other factors we did not model). A priori it was not obvious that human errors would be so closely replicated by an optimized system. Second, we used the model as an engine to screen a large set of spatial configurations, not all of which had been tested thoroughly in humans. This process yielded predictions of effects that we then confirmed in human experiments. Such cycles of human and model experiments illustrate the value of having stimulus-computable models of behavior.

### Limitations and future directions

Our modeling framework used a stereotyped task setting in which there is a cue stimulus from which attentional gains can be derived, followed by a stimulus to which the attentional gains are applied. This setting facilitated large-scale training and testing of attention, but does not fully capture the variety of ways in which real-world attention arises. Some of the additional complexity found in real-world attention likely reflects flexible executive control of attention. For instance, humans can direct auditory attention to a verbally instructed location, or to a familiar voice that comes to mind even in the absence of a prior cue stimulus. We envision that the same basic attentional gain mechanisms could work in these settings, but with the gains derived from an internal representation rather than the cue stimulus. Humans can also adjust the strength of attention using executive control, perhaps based on the perceived difficulty of a task, and potentially relating to the feeling of effort. Extending the modeling framework to allow the strength of attentional gains to vary could help to understand effort in computational terms.

Another complexity that we neglected is that attentional gains are likely refined over time, as the experience of attending to a particular sound source plausibly causes the representation of the source’s properties to become more precise, which could be profitably incorporated into the attentional “filter”. Evidence for an evolving attentional filter comes from findings that humans can use attention to track sources whose features change over time^85,86^. Accounting for these abilities would require a more complicated way to set attentional gains based on the selected source. The ability to refine attention over time may be particularly important when attention is based on an abstract memory of a class of target stimuli (e.g. the sound of a motor vehicle, or of a woman’s voice), rather than a specific recently heard stimulus. The abstract memory may serve as an initialization of attentional gains, that are then updated based on the experience of listening to the actual sound source encountered by the listener.

Another limitation of our model is that attention depends only on the cue stimulus. In some settings, attentional gains might additionally be shaped by the properties of stimuli to be ignored. Models whose gains are also a function of the mixture could be used to investigate this idea. Models like ours could also provide hypotheses for the representation of stimuli outside the focus of attention^1,17^, for instance by measuring what can be decoded about unattended stimuli from different model stages.

The models we built here are composed of simple operations that are loosely inspired by neuroscience, but deviate in many ways from biological sensory systems, making them inappropriate as models of some neural phenomena related to attention^87^. However, as it becomes possible to train models that are more biologically realistic^88^, the general framework we propose here should remain applicable. Lastly, our framework is modality-independent, and extensions to other modalities could help reveal whether general principles govern feature-based attention in vision and hearing.

## Methods

### Informed consent

All participants provided informed consent in accordance with the Massachusetts Institute of Technology Committee on the Use of Humans as Experimental Subjects. All participants received compensation for their involvement. No experiments were pre-registered.

### Training data generation

The training dataset consisted of 3,973,192 labeled exemplars, with each exemplar consisting of a cue signal and a multi-source mixture with a corresponding word label. Cue signals were two-second binaural audio clips of a single “target” talker spatialized to a location (defined by an azimuth and elevation) relative to a simulated listener position. Mixture signals were another two-second binaural audio clip: a superposition of a different excerpt of the target talker (rendered at the cued location) and other spatialized audio signals (either excerpts of other talkers and/or non-speech sound sources, rendered at locations that could be distinct from that of the target). Word labels corresponded to the word spoken by the target talker that overlapped the middle (i.e., the 1-second mark) of the mixture. Below, we will describe the curation of source materials included in training, the room simulator used to spatialize sources, and the scene generation procedure.

#### Speech corpora

Speech excerpts used to train models were sourced from the English-language training split of the 9^th^ release of the Common Voice speech corpora. This corpus was selected for its large number of unique talkers and variety of speech materials. We screened the corpus to obtain a curated list of speech excerpts that would support our attentional word recognition task. The purpose of the screening was to balance occurrence of single-word classes, the number of utterances per talker, and talker sex for both targets and distractors across examples contained in the training set.

First, word boundaries were extracted from the recordings and transcripts using a forced alignment procedure described in previous publications^89,90^. We then removed examples that did not have a 44,100 kHz sampling rate, helping to ensure that excerpts natively contained the frequency content implicated in supporting human sound localization^91^. Words spelled with less than five characters or spanning longer than two seconds in duration were excluded. From the remaining set of word excerpts, we took at most 5,000 examples per word to limit word class imbalance, randomly sampling from the available examples for word classes exceeding this limit. The top 800 most numerous words were taken to be our training vocabulary. Word class balance was then obtained by resampling each word class to have 5,000 exemplars, drawing from the screened set with replacement. This returned 3,994,484 million total single-word examples, with 2,041,825 unique utterances before up-sampling word classes.

Of the 15,735 unique talkers that remained in our training set after the prior screening steps, 3,407 were female talkers and 12,325 were male talkers. To avoid overrepresenting a particular talker sex when generating training scenes, our final screening step was to filter the 3,994,484 million excerpts to obtain a sex balance at the excerpt level. We did this by taking all available examples of female talkers in the word-balanced set (496,649 of 3,994,484), then sampling the same number of male-talker examples from the remaining set. The resulting screened set holds 993,298 total single-word examples (496,649 female, 496,649 male). Cue, target, and distractor speech excerpts were sampled from this screened set of single-word examples when constructing the final training set of cocktail party scenes.

Validation set examples were generated using the same screening procedure described above, sourcing materials from the corresponding validation split of Common Voice. For the validation set, we sampled up to 250 examples per word class instead of 5,000 as in training, resulting in 196,968 single-word excerpts. Gender balancing was not imposed for the validation set.

#### Noise corpora

Natural sounds clips were sourced from AudioSet^92^. We screened the entirety of AudioSet to select sound excerpts that were appropriate for our scene generation procedure. The main goal was a) to avoid speech content, as we wanted all speech in the dataset to come from the speech corpora described above, and b) to ensure that the sampling rate was high enough to be appropriate for spatialization. We first screened AudioSet examples to find a curated list of “parent” clips, then excerpted individual clips from these parent clips. We first removed AudioSet examples labeled with “Music”, “Speech”, “Singing”, “Vocal music”, “Whispering”, “Shout”, or “Silence”. As with the Common Voice excerpts, we then removed examples that did not have a 44,100 kHz native sampling rate. Parent clips were also screened to be at least 9-seconds in duration, enabling many 2.5-second excerpts per example. We performed the screening separately on the original training and validation split assignments of AudioSet. This returned 661,877 parent clips for training and 14,570 parent clips for validation. Individual natural sound clips were taken from these parent clips as part of the scene generation procedure. To obtain each natural sound clip in a scene, we first sampled a parent clip (uniformly from the screen training clips), then sampled a 2.5-second excerpt from the parent clip via a uniform random crop.

#### Virtual room simulation

To spatialize scenes, we used the same room simulator and a similar set of room simulations as Saddler and McDermott (2024)^22^. The description of the simulator is reproduced from their paper apart from minor edits indicating where our parameters differ from theirs. The simulator used the image-source method, incorporating KEMAR HRTFs, to render sets of binaural room impulse responses (BRIRs) for 2000 different shoebox-style rooms. Room dimensions were sampled log-uniformly between 3 and 30 m for length and width, and between 2.2 and 10 m for height. The listener’s head position was sampled uniformly in each room, under the constraint that the head was at least 1.45 m from every wall and no higher than 2 m from the floor. BRIRs for 1584 source locations (2 distances from the listener × 72 azimuths × 11 elevations) were rendered for each listener environment. One of the distances was 1.4 m for every BRIR. The other distance was independently sampled for each BRIR (drawn uniformly between 1 m and 0.1 m less than the distance from the listener to a wall). 1800 unique listener environments were included in the training set and the remaining 200 were used for validation. The final training and validation datasets consisted of 2,851,200 and 316,800 simulated positions, respectively. Our simulated spatialization differed from that in Saddler and McDermott (2024) only by the inclusion of negative elevations: we used 11 total elevations compared to the 7 in Saddler and McDermott, giving 1,584 source locations per listener environment with 2,851,000 total positions (vs. 1,008 source locations per listener environment and 1,814,400 total positions in Saddler and McDermott).

#### Training data scene generation

Auditory scenes were created by combining talker clips from the curated set of Common Voice examples with natural sound clips from the curated set of AudioSet examples, which were then spatialized using the room simulator. Scene parameters were varied across training exemplars to sample a wide range of conditions. To generate a training exemplar, we used the following procedure. First, we sampled a target excerpt from the curated list of Common Voice examples. Second, a cue excerpt was sampled from the set of clips produced by the target talker, restricted to be centered on a different word than the target clip. Third, we sampled the total number of distractor sources (uniformly from 1 to 6 sources). Fourth, we sampled the number of these distractor sources that were talkers (uniformly from 0 to n, with n being the sampled number of distractors for the scene). Fifth, we sampled the distractor talker clips from the Common Voice examples not produced by the target talker. Sixth, we sampled the remaining distractor sources from the curated set of AudioSet excerpts. Individual natural sound clips were taken from an AudioSet excerpt by first uniformly sampling a parent clip, then randomly excerpting a 2.5-second crop from that parent clip. Seventh, the sound pressure level of each distractor source was uniformly sampled from 50 dB SPL to 70dB SPL.

In each scene, the target and cue source were localized to the same azimuth and elevation, relative to the sampled listener position. To prevent the model from exclusively exploiting localization cues, half of the training scenes had distractor sources at the same location as the target source. For the remaining half of training examples, the azimuth and elevation of each distractor source were uniformly sampled from the possible source locations relative to the listener position. After setting distractor clip levels, distractor clips were spatialized to their selected locations then combined (summed in each channel).

To increase variability in the training data, each target clip in the curated set of Common Voice was sampled four times, each time being part of a different scene. The final composition of the training set contained 3,973,192 unique exemplars (1,988,317 female; 1,984,875 male). Validation scenes were generated via the same procedure, but with clips sampled from the validation set source materials, resulting in 196,968 exemplars.

#### Signal augmentations

Cue, target, and distractor-scene compositions were pre-sampled and stored separately, enabling augmentations to be applied to the target clips and distractor scenes. To prevent models from conflating the characteristics of the recording conditions for an individual talker with properties of their voice, bandpass filters were applied as augmentations to either the cue or target excerpt for 50% of the training examples. Filtering was performed using digital Butterworth filters, stochastically choosing filter parameters for each example, uniformly sampling low-frequency cutoffs from 40Hz to 400Hz, high-frequency cutoffs from 4kHz to 16kHz, and the filter order from 1 to 4. To avoid having models overfit to the onset times of labeled words, target clips were randomly shifted in time (either forward or backward with equal likelihood). Time shifts were uniformly sampled from 0 to 50% of the labeled word’s duration, constrained so the labeled word still overlapped the middle (1-second mark) of the target clip after shifting. Mixture clips were obtained online during training by superimposing the target and distractor scenes at a signal-to-noise ratio (SNR) uniformly sampled from −10 dB SNR to 10 dB SNR. Cue and mixture clips were then RMS (root-mean-square) normalized to 0.02. Finally, to allow models to learn how to report words said in isolated speech excerpts without being cued, 10% of training examples features single-talker target signals with silence as a cue. This was done by simply not mixing the target with the paired distractor and using silence (an array of zeros) instead of the paired cue signal for these examples.

#### Boundary handling

To avoid signal onset/offset artifacts, all cue, target, and mixture signals were extracted to be 2.5 seconds in duration. Once augmentations were applied, signals were passed through the cochlear model, after which the middle 2-seconds were excerpted as the final stimulus.

### Model implementation

Models were implemented in the PyTorch deep learning library using the PyTorch Lightning framework for efficient training in distributed settings. All model training and analysis used the compute resources of MIT’s OpenMind computing cluster, running Python version 3.12, Pytorch version 2.4, and Pytorch Lighting version 2.1. All Python dependencies and package versions are available in the code repository that will be made available upon publication.

### Cochlear model stage

The first stage of our models was a fixed simulation of the cochlea and auditory nerve, providing a ear-by-time-by-frequency representation intended to replicate the auditory cues provided by the human auditory periphery. This initial stage took the sound waveform as input. The input was passed through a finite-impulse-response (FIR) approximation of a gammatone filter bank (with impulse responses truncated to 25 ms to reduce memory consumption) the output of which was half-wave rectified, passed through a compressive nonlinearity, then downsampled. These operations were performed separately on the left and right audio channels.

First, each of the two 44.1kHz stereo audio waveforms was convolved in the time domain with the FIR approximation of a 40-channel gammatone filter bank (1,102 taps per filter) with center frequencies spaced uniformly on an ERB-numbered scale between 40Hz to 20kHz. Second, the resulting subbands were half-wave rectified. Third, the half-wave rectified subbands were raised to the power of 0.3 to simulate the compressive response mediated by outer hair cells. Fourth, the compressed, rectified subbands were low-pass filtered with a 4kHz cutoff and down-sampled to 10kHz, to both impose the upper limit on phase locking of inner hair-cells and reduce the dimensionality of the neural network inputs. Low-pass filtering and down sampling was performed via 1d convolution with a Kaiser-windowed sinc filter, with a filter width of 64, roll-off of 0.94759, and beta of 14.76965, implemented with the Torchaudio transforms resample method. Finally, to avoid signal onset/offset artifacts, the middle 2-seconds were then excerpted from the full 2.5-second input signal duration. We refer to the resulting representation as a “cochleagram”. The left and right cochleagrams were concatenated, yielding a 2-channel by 40-frequency by 20,000-timestep input to the neural networks.

### Neural network model stages

Models of feature-based attention were built using a convolutional neural network backbone equipped with feature-based gains as parameterized sigmoid functions. The neural network backbone consisted of a cascade of convolution, pooling, and normalization operations, as in previous models from our lab^22,30,37,93,94^ which have yielded strong task performance and close matches to human behavior on other auditory tasks, as well as state-of-the-art predictions of brain responses from human auditory cortex^30,95^. Below, we will detail the core operations included in our artificial neural network architectures, the implementation of feature-based gain, the forward-pass used in our training algorithm, the architecture search process used to select model architectures, and the modifications made to constrain attention in our control models.

### Feature-based gain implementation

Feature-based attention was implemented as sigmoidal functions with learnable parameters at each neural network model stage:

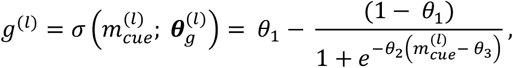

where (*l*) indicates the network stage, ***θ***_*g*_ = [*θ*_1_, *θ*_2_, *θ*_3_] are learned parameters for the bias, slope, and threshold of the function, 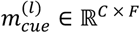 is the time-averaged representation of the cue at layer (*l*) (of size C channels by F frequencies), and *g*^(*l*)^ ∈ ℝ^*C* × *F*^ are the feature-gains for each of the *C* × *F* features. Gain functions were applied to the cochleagram and to each convolutional block in the neural network.

### Forward pass through the feature-gain models

The forward pass for our feature-gain models augmented the standard forward pass used in feed-forward networks, enabling stage-specific application of gains derived from the cue representation at a given stage to the mixture representations at the same stage. Given a cue *x*_*cue*_ and mixture *x*_*mix*_, the forward pass of the model runs as follows. First, we obtain a representation of the cue for each model stage

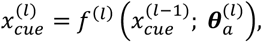

where 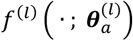 *i*s the *l*_*th*_ model stage (i.e., convolutional block), 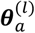 are the parameters for that model stage, and 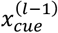 is the cue representation from the previous model stage. Second, a memory representation, 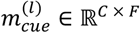, of the cue, 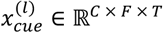, is obtained by averaging over the time dimension of the cue representation at the same model stage:

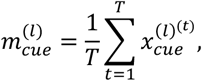

Third, gains are then obtained from the memory representation

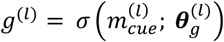

where 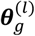 are parameters for the gain function at stage *l*. Fourth, the mixture representation is obtained

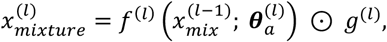

where 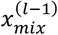 is the mixture representation from the previous stage, 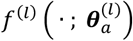 is output of the *l*_*th*_ convolutional block, *g*^(*l*)^ are the feature-gains obtained from the cue, and ⊙ is the element-wise multiplication operator. For simplicity, the same set of feature-gains were applied to all timepoints of the corresponding mixture representation. After the final convolutional stage, the mixture representation was passed through a fully-connected layer followed by the 800-dimension linear output stage for the word recognition task.

During this forward pass, the weights of the convolutional blocks, ***θ***_*a*_, are shared when obtaining cue and mixture representations for each training example.

### Artificial neural network architectures

The operations in the convolutional neural network architectures were organized as a series of convolutional blocks. Each convolutional block consisted of a sequence of layer-normalization, convolution, point-wise nonlinearity, and pooling operations. The final convolutional block of each architecture was followed by a sequence of four operations: a single fully-connected layer, a point-wise nonlinearity, dropout, and a softmax classifier.

#### Layer normalization

Normalizing activations between neural network stages improves the efficiency of ANN training by helping to stabilize gradient updates: the magnitude of a parameter update in one stage is less likely to be amplified in following stages if they are separately normalized. Normalization-like operations are also common in biological sensory systems, and have been proposed to interact with multiplicative gains to produce attentional effects observed neurophysiologically^14^. Layer normalization^96^, a common choice for normalization operations in ANNs, point-wise normalizes input examples individually using the mean and variance over feature dimensions, with high output reflecting high activity relative to the features of that input example. We used layer normalization (as opposed to batch normalization^97^, the other common choice for ANN normalization operations) because it is more similar to the normalization found in sensory systems in normalizing responses by a function of the current stimulus^98^, rather than by a function of the training distribution. Layer normalization also does not require scaling test examples to match the statistics of the training examples (as is necessary with batch normalization), which might aid generalization at inference. The layer normalization operation is defined as

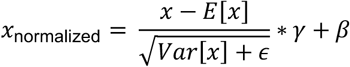

where *x* ∈ ℝ ^*C* × *f* × *T*^ is the input tensor, *E*[*x*] and *var*[*x*] are the mean and variance over feature dimensions (*C, f, T*) of *x, ϵ* = 0.00001 to prevent division by zero, and *γ* ∈ ℝ ^*C* × *f* × *T*^ and *β* ∈ ℝ ^*C* × *f* × *T*^ are tensors of learnable parameters.

#### Convolution

Each convolutional layer consisted of a bank of learnable filter kernels 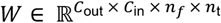, with *C*_out_ different kernels, *C*_in_ input channels, and kernel dimensions of *n*_f_ by *n*_t_ in frequency and time, respectively. Inputs to each convolutional layer, 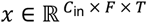, were three-dimensional tensors with *C*_in_ input channels, *f* features, and *T* time samples. For the first convolutional layer, *f* = 40 and *T* = 20,000 (the frequency and time dimensions of the cochleagram), while *C*_in_ = 2 (the left and right audio channels).

Boundary handling for convolution operations was identical to that in Francl and McDermott (2022). Specifically, “valid” convolution was used in the time dimension (i.e. no zero-padding applied), and “same” convolution was used in the frequency dimension. The rationale for this choice was to avoid temporal edge artifacts that would otherwise result from zero-padding in the time dimension. Edge effects in the frequency dimension are less clearly inconsistent with biology because the cochlea has upper and lower frequency limits, and were considered preferable to the rapid loss of dimensionality along the frequency axis that would occur with “valid” convolution given the small number of input frequency channels.

For an input tensor *x*, the output of a convolutional layer is a tensor 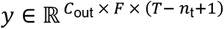 given by:

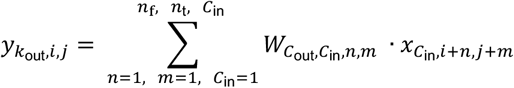

The output from each convolution thus had dimension (*C*_out_ × *f* × *T* − *n*_t_ + 1). Because convolutional layers were preceded by layer-normalization, bias vectors were omitted (to minimize redundant parameters and reduce memory consumption). All convolutional layers used a stride length of 1.

#### Point-wise nonlinearity

Nonlinear activation functions enable neural networks to learn complex functions. We used rectified linear units as the point-wise nonlinearity in all architectures. This operation is defined as

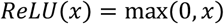

#### Weighted-average pooling

Pooling layers downsample their inputs by aggregating information across neighboring frequency and time points. Weighted average pooling with Hanning windows was used to reduce aliasing in our networks (which would occur if downsampling was not preceded by lowpass-filtering)^99^.

Pooling was performed by convolving input tensors *x* ∈ ℝ ^*C* × *f* × *T*^ with two-dimensional (frequency by time) Hanning window kernels,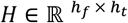:

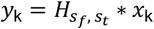

where * is the convolution operation, k indexes the channel dimension, and *s*_*f*_, *s*_*t*_ are the stride length in frequency and time, respectively. As in Saddler et al., (2021)^93^, the Hanning kernel had a stride-dependent shape, where

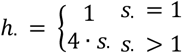

for *h*_*f*_ and *h*_*t*_ depending on *s*_*f*_ and *s*_*t*_, respectively. For an input *x* ∈ ℝ ^*C* × *f* × *T*^, the corresponding output is 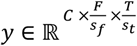. If either *s*_*f*_ or *s*_*t*_ equal 1, no pooling is performed on the corresponding dimension.

#### Fully-connected layer

A fully-connected (also sometimes called linear or dense) layer applies an affine transformation to an input tensor *x* ∈ ℝ ^*D*^. In our networks, where *x* ∈ ℝ ^*C* × *f* × *T*^ is the output of a convolutional block, *x* is first flattened to a vector *x*_flat_ ∈ ℝ ^*D*^ where *D* = *C* · *f* · *T*. Then, an affine transform is applied

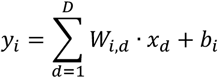

producing an output vector *y* ∈ ℝ ^*N*^, where *W* ∈ ℝ ^*N* × *D*^ are learned weights, and *b* ∈ ℝ ^*N*^is a learned bias vector.

#### Dropout regularization

Dropout is a form of regularization applied to input tensors *x* ∈ ℝ ^*N*^ during training, intended to minimize co-adaptation of units. On each forward pass, a fraction *P* of the units in *x* are sampled (from a Bernoulli distribution) and set to zero. The remaining fraction of units are scaled by 1/(1 − *P*) so the expected sum over all outputs is the same as the expected sum over *x*. During inference, the operation is replaced by the identity function. Dropout was applied during training to the activations of the penultimate fully-connected layer preceding the softmax classifier in our networks, with *P* = 0.5.

#### Softmax classifier

A softmax classifier was the final layer in all our networks. The softmax classifier is a composition of a fully-connected layer and a normalized exponential function. First, input tensors *x* ∈ ℝ ^*D*^ are passed through a fully-connected layer *h*( ·, *W*), with weights *W* ∈ ℝ ^*V* × *D*^ mapping from *D* features to *V* word classes. The operation returns un-normalized activations for each word class (often called logits):

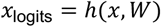

These logits are then scaled by the softmax function producing an output vector *y* ∈ ℝ ^*V*^:

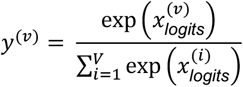

where *v* indexes the word class. Because the values of *y* are greater than zero and sum to one, *y* can be interpreted as a probability distribution over word classes for the given input mixture.

### Architecture selection

Artificial neural network performance depends both on the network weights learned during gradient descent, and the configuration of the operations (defined by “hyperparameters”) that define an architecture (i.e. the number of layers, number of channels per layer, and the operations in each layer). To ensure we used optimal hyperparameter settings in our CNN backbone, we drew 13 architectures from successful models of word recognition and sound localization identified by prior work^22,30,37,93,94^. First, we piloted all training and model experiments using one of these architectures (the top architecture in Saddler & McDermott, 2024^22^), which yielded strong matches to human behavior. Then, we trained the remaining 12 and selected the top 10 architectures based on their validation set performance. Model results for each experiment are reported as the average across these 10 best network architectures, enabling us to report measures of uncertainty and marginalize across the eccentricities of any single network architecture. Hyperparameters of these 10 architectures are in Supplementary Table 1.

### Models with alternative architectural constraints

To test whether explicit feature gains were necessary for attentional selection, we instantiated models that altered how gains were included in the model architecture. We tested three control architectures that each informed a particular hypothesis. The first was a “gain-free” architecture, that tested the necessity of including explicit gain functions. The second was a model with gains only at the early stage of the architecture, that tested whether selection of low-level features was sufficient for task performance. The third control model included gains only at the late stage of the architecture, testing whether selection of higher-level features was sufficient for task performance. Each control architecture used the backbone CNN of the best-performing feature-gain model (architecture 1 in Table 1). Control models were trained using the same training set and optimization hyper-parameters as the feature-gain architectures. The details of each control architecture are given below.

Baseline model without explicit feature-based gains: We trained a version of architecture 1 using the backbone CNN without gain functions. Instead, cue and mixture signals were concatenated along the channel dimension and passed as a single input *x* ∈ ℝ^*C x f x T*^with dimensions *C* = 4 channels (the stereo channels for cue and mixture), *f* = 40, and *T* = 20,000 (the frequency and time dimensions of each cochleagram). The model architecture was therefore augmented to accept 4 input channels (compared to 2 as in the feature-gain models), using the forward pass of a traditional CNN. This model tests the extent parameters of the CNN architecture alone support solutions to the attentional selection task, providing a performance baseline models with feature gains can be compared to.

Model with feature-based gains only at early stages: Inputs to the model, the model forward pass, and application of gains were identical to the unaltered feature-gain models, with the exception that gains were only applied to the cochlear representation of the mixture before the first convolutional layer. This model tests the hypothesis that constraining selection to low-level features available in the early auditory pathway is sufficient for enabling attentional selection.

Model with feature-based gains only at late stages: We modified architecture 1 to include attentional gains only between the last convolutional layer and the fully-connected layer. Model inputs, the model forward pass, and application of gains were identical to the full feature-gain models, with the exception that gains were only applied to the outputs of the final convolutional layer. Consequentially, this model tests whether selection of features from higher-order representations alone is sufficient for enabling attentional selection.

Model with alternative location of feature-based gains: This model architecture was identical to the main feature-gain architecture except that the gains were applied after the normalization operation rather than before it.

### Model optimization

All models described above were trained using the dataset described above via stochastic gradient descent, using the AdamW^100^ optimizer (with a learning rate of 0.00005 and a batch size of 288). Models were trained using a distributed data parallel strategy run on 4 NVIDIA A100 GPUs, where each GPU ran a unique subset of a training batch through a copy of the model parameters (which were updated synchronously with respect to the whole-batch loss) enabling larger batch sizes in training. 16 CPUs (4 per GPU) and a total of 100Gb of memory were used to execute data reading and online signal augmentations during training for each model. All models were trained until performance on the validation set converged (approximately 10 training epochs of the training set).

### Human behavioral experiments

Human experiments were conducted both online and in-person, depending on the type of experiment. All participants in both online and in-person experiments self-reported as native English speakers having no known hearing loss.

All diotic listening experiments (Experiments 1-3) were conducted online to facilitate large sample sizes (to increase the reliability and reproducibility of the results). Extensive research conducted in our laboratory has consistently demonstrated that online data can match the quality of data collected in traditional laboratory settings^30,101-106^ provided measures are taken to ensure standardized sound presentation, encourage participant compliance, and promote active engagement in the tasks. Online participants were recruited using the Prolific platform. To ensure data quality, participants were prescreened to have at least a 95% submission approval rate, and to have not completed any online experiment hosted by the authors’ lab in the past 6 months (to avoid familiarity with our experimental stimuli and task). All online participants were instructed to complete the experiment in a quiet environment and completed a headphone check experiment^107^ prior to the main experiment to help maximize sound presentation quality. Participants adjusted the volume of a calibration sound to a comfortable level at the start of the experimental session, and all stimuli were scaled relative to this maximum level. Participants also completed 12 catch trials in each experiment. These were intended to make sure participants were paying attention to the experiment and served as an additional screening metric. Catch trials presented isolated words (spoken by one of the authors) in silence using the same clip for cue and mixture and were randomly intermixed with experimental trials. Participants scoring less than 11 correct catch trials (91.6% accuracy) were excluded from our analysis.

Experiments 6 and 7 measuring the effects of spatial separation on attentional selection were run in-person over a speaker array (detailed below) to ensure the accuracy of stimuli playback location. Because participants were monitored for the duration of these experiment, catch trials were not included.

### Experiment 1: effect of distractors on attentional selection in monaural conditions

Experiment 1 measured cued word recognition as a function of signal-to-noise ratio (SNR) and distractor type. Talkers and speech materials were not included in model training so as to enable fair comparisons between models and humans. Nine distractor types were used: 1-talker same-sex, 1-talker different-sex, 2-talker, 4-talker, 8-talker babble, 1-talker Mandarin speech, stationary noise, recorded auditory scenes, and instrumental music.

#### Stimuli

Cue, target, and English-speech distractor signals were sourced from the portion of Spoken Wikipedia Corpora^108^ material screened in Feather et al^99^. Feather et al. removed Spoken Wikipedia articles due to a) potentially offensive content for human listening experiments, b) missing data or c) having bad audio quality (for example, due to computer generated voices of speakers reading the article or the talker changing mid-way through the clip). We applied further screening to this set to identify clips centered on words in the model vocabulary (316,748 clips), so that the model could also be run on the experiments.

Target clips were additionally constrained to be sex-balanced. The in-vocabulary clips were further screened to find words with at least one example produced by both a male and female talker, yielding 488 target words. One male- and one female-talker example was uniformly sampled for each target word, giving 976 total target clips. Cue clips were drawn from target excerpts centered on words not included in the model vocabulary.

English speech clips for the 1-, 2-, and 4-talker distractor conditions were sampled from the full set of Spoken Wikipedia excerpts containing words in the model vocabulary. This made it possible to measure the rate of distractor word reports without needing to reuse target clips as distractors in a given experiment. So that the effect of talker-sex similarity could be analyzed in the 1-talker distractor condition, one male and one female distractor clip were sampled for each target clip. Two 2-talker distractor clips were sampled for each target clip by summing either two male or two female clips. The 4-talker distractor clips were created by summing the two male and two female distractor clips. The talker identity and middle word of the distractor clip(s) were constrained to differ from those of the target clip. The 1-talker Mandarin distractor clips were sourced from the Mandarin validation (termed “dev” in the dataset documentation) portion of Common Voice (version 9). One male and one female Mandarin speech clip were sampled for each target clip to be included in the talker sex analysis. Speech-shaped noise was synthesized for each target clip by imposing the power spectrum of the target clip on white noise. Instrumental music, auditory scenes, and 8-talker babble were sampled from the MUSDB18, IEEE AASP CASA Challenge, and Common Voice test clips used in Saddler & McDermott (2024) (450, 400, 400 total clips respectively).

Target clips were combined with distractor clips from each condition at 6 SNRs (infinite, i.e. no-distractor, and −9, −6, −3, 0, +3 dB) producing 44,896 possible mixture stimuli (976 target clips with no distractor + 976 target clips x 9 distractor types x 5 SNRs). Cue, target, and distractor signals were 2-seconds in duration, sampled at 44,1 kHz. Mixtures were obtained by setting the target clip against the distractor clip at the given SNR. All cue and mixture clips were normalized to a RMS of 0.02 for model experiments, and were presented at the same participant-determined level for the human experiments (described in the Procedure section).

#### Procedure

Individual participants completed 12 catch trials and 184 experimental trials (4 trials x 5 SNRs x 9 distractor conditions + 4 no-distractor trials). Each participant heard a random sample of 184 of the 488 target words, with the talker sex randomly determined for each word. The words were then randomly assigned to distractor and SNR conditions for each participant, constrained to yield both a sex balance over target talkers and target-distractor sex conditions, and unique cue and mixture clips. Target-distractor pairings were constrained as described above. The 12 catch trials were randomly intermixed with the 184 experimental trials.

On every trial, participants first heard the 2-second cue, then a 0.5-second delay, then the 2-second mixture. This audio sequence was initiated by a button click. Participants reported the word said by the target voice in the middle of the mixture clip (defined as the word overlapping the 1-second mark of the mixture). To equate the task for humans and models, participants were asked to select words from a list of the 800 words in the model vocabulary. Participants typed responses into a text box and, while they typed, the web page displayed a list of 800 words that was continuously updated to only include matches to the string they typed. Only words in the word list could be entered, analogous to how the models could only report one of these 800 words. Participants received feedback after each trial, displaying the trial outcome (“correct” or “incorrect”), the number of remaining trials, and their running accuracy. On incorrect trials, the correct word was also displayed to help participants lean to perform the task.

The experiment was run online over the Prolific platform. Participants first heard a calibration sound and adjusted the presentation level of their computer to be comfortable prior to the start of the experiment, and then completed a headphone check^107^ to help ensure good sound quality. Only participants who passed the headphone check completed the main experiment. We then only analyzed data from participants who responded correctly to at least 91% of catch trials and who self-reported normal hearing. 195 participants (98 female, 92 male, 4 non-binary, 1 no-report) met these criteria. Participants ages were between 18 and 71 (median 33) years.

#### Model experiment

Models were tested on all combinations of the 976 target clips (both male and female examples of each target word) with the 5 SNRs, 9 distractor conditions, and no-distractor condition (44,896 total stimuli) used in the human experiments. Because listeners used headphones rather than free-field speakers, audio signals given as input to the model were not run through the room simulator. Instead, each mono clip was presented diotically (i.e., the mono stimulus waveform was input for both the left and right channel of the model).

#### Analysis

To account for the possibility that participants correctly recognized the target speech signal but inadvertently reported a word that was adjacent to the trial’s target word rather than the target word itself, word recognition performance was measured by contrasting a participant’s response against all in-vocabulary words contained in a trial clip. In all conditions, a response was counted as correct if it matched any of the in-vocabulary words that were present in the target utterance. In the 1-distractor condition, confusions were analogously measured as reports of words contained in the transcript of the distractor utterance. This scoring was applied to both participant and model reports.

To analyze the effects of target-distractor sex similarity and language familiarity, we reanalyzed both the English and Mandarin 1-distractor conditions for both participants and models. To analyze target-distractor sex similarity, trials from both distractor language conditions were first pooled, then the mean for each distractor sex condition was obtained for each participant (or model architecture). To analyze the effect of distractor language, trials from both distractor sex conditions were pooled within language, then the mean for each distractor language condition was obtained for each participant (or model architecture).

### Experiment 1b: effect of cue duration (Supplementary figure 3)

It seemed possible that the fixed duration of the cue signal in our main task could limit the ecological validity of our results. To assess whether human and model behavior were robust to variation in the cue duration, we ran an additional experiment using the materials and procedure of Experiment 1, but with cues that varied in duration. The same set of cue, target, and single-distractor excerpts used in Experiment 1 were used, but cue signals were center cropped to be either 0.5-, 1-, or 2-seconds in duration. All target-distractor mixtures were presented at 0dB SNR.

#### Procedure

The procedure was identical to that of Experiment 1, with the following exceptions. Participants each completed 12 catch trials and 30 experimental trials (10 trials x 3 cue duration conditions). Each participant heard a random sample of 30 of the 488 target words, with the cue duration for each example randomly assigned for each participant. The 12 catch trials were randomly intermixed with the 30 experimental trials. 84 participants (43 female, 38 male, 3 non-binary) met our inclusion criteria (scoring 91% on the catch trials). Participant ages were between 20 and 40 (median 32) years.

#### Model experiment

Models were tested on all combinations of the 976 target clips (both male and female examples of each of the 488 target words) with the 3 cue duration conditions, and both distractor sex conditions (5,856 total stimuli) used in the human experiments. As in Experiment 1, the mono audio clips were not run through a room simulator, and were instead presented to the model diotically.

### Experiment 2: effect of harmonicity on attentional selection in monaural conditions

Experiment 2 measured how violations of harmonicity (the tendency of frequency components to be integer multiples of a fundamental frequency) impact cued word recognition. Participants were presented with a cue, followed by either the target excerpt alone, or the target excerpt mixed with a distractor excerpt. Harmonic, inharmonic, and whispered stimulus variants were generated as in Popham et al., 2018 (summarized below), using the stimuli from the English 1-distractor condition of Experiment 1. Twelve target-distractor harmonicity conditions were used (3 target harmonicities – harmonic, inharmonic, and whispered – x 3 distractor harmonicities + 3 target harmonicities with no distractor). We only analyzed the conditions in which the target and distractor were of the same type (harmonic/harmonic, inharmonic/inharmonic, and whispered/whispered), to be consistent with the original experiment run by Popham et al. Participant and model responses were analyzed identically to Experiment 1.

#### Stimuli

Experiment 2 used the set of cue, target, and distractor pairings from the 1-talker same-sex and 1-talker different-sex conditions of Experiment 1. All stimuli were resynthesized from the original speech excerpts of Experiment 1 using the STRAIGHT algorithm^109,110^. Inharmonic speech examples were synthesized by shifting harmonic frequency components above the fundamental frequency by either −30% or 30% (which maximally reduced intelligibility in Popham et al., 2018). Jitter values were sampled independently for each harmonic’s frequency of each original speech clip but were constrained (via rejection sampling) such that adjacent harmonics were always separated by at least 30 Hz. To generate the stimuli for an inharmonic trial, one jitter pattern was sampled for the target excerpt, and this same jitter pattern was applied to the cue and distractor utterances. Harmonic stimuli were synthesized by running stimuli through STRAIGHT without changing any of the constituent harmonic frequencies. Whispered speech was created as in Popham et al. 2018, and the description of the synthesis from that paper is reproduced here with minor edits for clarity. Whispered stimuli were generated by omitting the sinusoidal excitation used for harmonic/inharmonic synthesis and high-pass filtering the noise excitation to simulate breath noise in whispered speech. The filter was a second-order high-pass Butterworth filter with a (3 dB) cutoff at 1200 Hz whose zeros were moved toward the origin (in the z-plane) by 5%. The resulting filter produced noise that was 3 dB down at 1600 Hz, 10 dB down at 1000 Hz, and 40 dB down at 100 Hz, which to the authors sounded like a good approximation to whispering. Without the zero adjustment, the filter removed too much energy at the very bottom of the spectrum. The noise excitation was combined with the time-varying spectral envelope using the same procedure employed for harmonic and inharmonic speech. The noise-excited stimuli were thus generated from the same spectrotemporal envelope used for harmonic and inharmonic speech, just with a different excitation signal.

Experiment 2 used the same combinations of cue, target, and distractor source clips as Experient A. Harmonic, inharmonic, and whispered versions of each target-distractor pairing were synthesized as described above. To produce the 9 target-distractor harmonicity combinations, each version of a target clip was crossed with each version of the paired distractor clip. Cue clips always used the harmonicity of the target clip. Signal sampling rates and presentation levels matched those of Experiment 1.

#### Procedure

The procedure of Experiment 2 was identical to that of Experiment 1, with the following exceptions. Participants each completed 12 catch trials and 180 experimental trials (15 trials x 12 harmonicity conditions). Each participant heard a random sample of 180 of the 488 target words, with the harmonicity condition for each word randomly assigned for each participant. The 12 catch trials were randomly intermixed with the 180 experimental trials. 90 participants (43 female, 47 male) met our inclusion criteria (scoring 91% on the catch trials). Participant ages were between 19 and 64 (median 34.5) years. We note that this experiment used a different type of cueing than was used in the original experiment of Popham et al. (the Popham et al. experiment showed participants the first two words of the sentence they were supposed to listen to, prior to the start of the audio stimulus). This task difference may be responsible for the lower rate of confusions we observed compared to the Popham et al. experiment.

#### Model experiment

Models were tested on all combinations of the 976 target clips (both male and female examples of each of the 488 target words) with the 9 harmonicity conditions, and 3 no-distractor conditions (23,424 total stimuli) used in the human experiments. As in Experiment 1, the mono audio clips were not run through a room simulator, and were instead presented to the model diotically.

### Experiment 3: word recognition in naturalistic auditory textures

To obtain a stronger test of the extent to which the models captured human speech-in-noise perception, we ran our models on an experiment from Saddler and McDermott 2024^22^, which probed human speech-in-noise recognition across a large set of naturalistic distractor signals. The description of the human experiment is reproduced from the original paper with minor edits for brevity.

Human word recognition accuracy was measured in 43 different naturalistic auditory textures. 376 speech excerpts from the evaluation portion of the Word-Speaker-Noise dataset^99^ were embedded in 376 unique exemplars of each auditory texture. The 2s texture exemplars were previously generated^37^ to match the statistics of 43 recorded real-world textures^111^. The success of the iterative synthesis algorithm^111^ used to generate the textures was determined both subjectively (synthesized exemplars sounded like the recorded textures) and objectively (mean-squared errors between synthetic and original texture statistics were at least 40 dB below the mean-squared texture statistics of the original recordings). Speech excerpts were randomly assigned to one of the 43 texture conditions with the SNR fixed at −3 dB (each participant heard a different random assignment). The experiment was run online and included 47 participants (24 female, 23 male) who self-reported normal hearing, passed a headphone check^107^, completed at least 100 trials, and responded correctly to at least 85% of catch trials (isolated words presented in silence). Participant ages were between 23 and 59 (median 39) years old.

#### Model experiment

We simulated a version of the original human experiment, modified to make the task compatible with the model task. For each target excerpt, a corresponding cue was obtained by sourcing a different excerpt of the target talker from the Word-Speaker-Noise dataset. We then measured model word recognition accuracy for speech embedded in each of 43 auditory textures, using the same stimuli as in the human experiment. Models were evaluated on the full stimulus set (16,168 stimuli = 376 speech excerpts × 43 auditory textures).

### Experiment 4: spatial tuning for masked speech

We simulated experiment 1 of Byrne et al. (2023)^34^, which measured spatial tuning for the release of masking of target speech, manipulating the distractor type and proximity to the target source. The original experiment used a closed-set masked speech identification task^112^, in which listeners were presented with 5-word sentences, in English with fixed syntactic structure, spoken by one of 12 female talkers (stimuli fully described in Kidd et al.^112^). Listeners were cued to the target stream by being told the starting word of the target sentence, which was fixed throughout the experiment, and that the target would occur directly in front of their position. Three distractor conditions were used: 1) two competing female talkers drawn from the 11 remaining talkers, 2) time-reversed speech made from the same sources used in condition 1, and 3) speech-shaped speech-envelope-modulated noise derived from the speech distractors of condition 1. The target source was randomly selected per trial, and was always located at 0° elevation and 0° azimuth. Two distractor sources were positioned symmetrically in azimuth around the target talker at angles of ±0°, ±5°, ±10°, ±20°, and ±40° azimuth and 0° elevation. For each distractor x angle condition, speech reception thresholds were measured using a one-down-one-up adaptive procedure, varying the signal-to-noise ratio between the target and summed distractor signals to estimate the 50% correct point on the psychometric function. The distractor level was held constant such that the sum of the two distractors was 60 dB, and the target level was adjusted by 3dB increments depending on participant accuracy. Trials were evaluated as correct, and the target level was reduced, if participants correctly identified the remaining 4 words in the target sentence. We reproduced the first of the three conditions (the one with two competing talkers) in our model experiment. Data was collected for 18 normal hearing native English speaking listeners (11 female, 6 male, 1 non-specified; aged 18-40 years). We scanned the results in from the original paper figure.

#### Model experiment

We simulated a version of the original human experiment using a virtual anechoic room and the same-sex distractor speech materials from Experiment 7. The model simulation differed in two respects from the human experiment. First, the target voice was designated by cue that was a different excerpt of the target talker (as in the other model experiments in this paper), rather than by one of the words spoken by the target talker. Second, in lieu of the adaptive procedure, we ran the model on 9 signal-to-noise ratios (−18 dB to 6 dB SNR in 3 dB steps) at each spatial location. All combinations of 976 stimuli (cue, target, and 1-talker same-sex-distractor pairs), 5 loudspeaker presentation conditions (±0°, ±5°, ±10°, ±20°, and ±40° azimuth), and 9 SNRs were simulated (43,920 total stimuli). Model thresholds for each loudspeaker condition were estimated as follows. First, word recognition performance was measured at each SNR and presentation condition.

Second, thresholds at each presentation condition were estimated by fitting a second-order polynomial to the performance by SNR curve, and calculating the SNR granting 50% of the model’s maximum performance.

### Experiment 5: the precedence effect with concurrent speech signals

We simulated experiment 1 of Freyman et al. (1999)^38^, which measured the benefit of perceived spatial separation on speech intelligibility, using the precedence effect to induce illusory spatial separation between concurrent talkers. The “precedence effect” traditionally refers to a perceptual phenomenon that occurs when two sounds played in quick succession from different locations seem to originate from the first location^35,36^. The effect has been hypothesized to reflect a strategy for localizing accurately in the presence of reflections. Because reflections typically arrive from a direction that is different from the direction of the source from the listener, they provide erroneous localization cues. However, they are also delayed relative to the direct sound from the source (which has cues that are faithful to the source location), because the path length traversed by the reflection is longer than that for the direct sound. Suppressing location cues from delayed components of a sound could thus make localization more robust to reflections. Consistent with this explanation, models that are optimized for localization in environments with reflections exhibit the precedence effect, whereas those optimized in anechoic environments do not^37^. Models plausibly learn to accentuate localization cues from early parts of a sound and suppress cues that are likely to originate from reflections, akin to effects seen in the early auditory system^39^.

In the experiment, a target talker was superimposed with a distractor talker. The experiment included a condition where the target and distractor were co-located and a condition where they were spatially separated, as well as conditions where the locations were illusorily shifted by adding a second copy of one of the signals at a different location, with a small temporal offset.

In the original experiment, 56 participants (sexes not reported) listened to recordings of speech, with one target talker and one distractor, and reported the keywords spoken by the target talker. Speech materials were of syntactically correct sentences that had no meaning. The target talker was always the same female voice, while the distractor was always the same alternate female voice. The experiment took place in an anechoic chamber, with loudspeakers positioned in a semicircular arc at 0° and 60° to the right relative to the listener position, at a distance of 1.9 meters from the listener. Six conditions were used for target and distractor locations (numbers reflect ordering in the original paper): 1) both target and distractor presented at 0°, 2) target presented at 0° with distractor presented at 60°, 3) target presented at 0° with distractor illusorily presented at 0° via precedence effect, 4) target presented at 0° with distractor illusorily presented at 60° via precedence effect, 5) both target and distractor illusorily presented at 0° via precedence effect, and 6) target illusorily presented at 0° and distractor illusorily presented at 60°. Illusory separation in conditions 3-6 was enabled by presenting signals from both positions and starting playback at the “perceived” location 4-ms before starting playback at the “lag” location (taking advantage of the precedence effect). Four signal-to-noise ratios (SNR) were presented per loudspeaker condition: −12 dB, −8 dB, −4 dB, and 0 dB SNR. We reproduce conditions 1, 2, and 4.

#### Model experiment

We simulated the original human experiment using a virtual anechoic room and the speech materials from Experiment 7. All combinations of 1,952 stimuli (cue, target, and single-distractor pairs), 3 loudspeaker presentation conditions (conditions 1, 2, and 4 described above), and the four original SNRs (−12 dB, −8 dB, −4 dB, and 0 dB SNR) were simulated (23,436 total stimuli). To simulate the precedence effect, we first spatialized the distractor to the lead channel. Next, we applied the 4-ms onset delay (by zero-padding) to the distractor, then spatialized it to the “lag” location. Finally, the two binaural waveforms were summed in each channel.

### Experiment 6: effect of spatial separation in azimuth and elevation

The benefit of target-distractor spatial separation was measured for separation in azimuth and elevation. Stimuli in Experiment 6 were a subset of the speech clips used in Experiments 1 and 3 that were screened to be suitable for sound localization experiments (described below). Experiment 6 was conducted in person, and stimuli were presented over a loudspeaker array (described below). Participant and model responses were analyzed as in Experiment 1

#### Stimuli

Speech excerpts for Experiments 6 and 7 were sourced from the set of Spoken Wikipedia Corpus examples used in the online monaural experiments (Experiments 1 and 3). Cue, target, and distractor stimuli were screened to include excerpts with frequency content beyond 16kHz to help enable sound localization in elevation. Target excerpts were further screened under the constraint that each word was spoken by both a male and female talker. These screening steps reduced the number of unique target words to 488 from 800, (976 total target excerpts, 488 per talker sex). This vocabulary size was large enough to allow for one unique word per trial in the in-person experiment. Cue excerpts were selected for each target excerpt by finding clips of the target talker centered on words not in the target vocabulary. To allow for analysis of talker-distractor sex similarity effects, each target excerpt was paired with both a same-sex and different-sex distractor. To enable use of symmetrically positioned distractors in the azimuth conditions, two distractors per sex condition were sampled from the screened set of distractors. This screening resulted in 1,952 unique combinations of cue, target, and two-talker distractors that were sex-balanced across both target talkers and target-distractor sex pairings.

#### Speaker array

All experiments measuring effects of spatial separation were run using a 19-by-5 array of loudspeakers arranged on a hemisphere (2 meter radius). Participants were seated at the center. Relative to the participant’s head, the array spanned 180° in azimuth (frontal hemifield) and −20° to 40° in elevation, with speakers spaced every 10° in both azimuth and elevation. Speaker locations were coded using labels that were attached to the bottom of each loudspeaker. Labels were formatted as a combination of one letter followed by two digits (e.g. A10), with letters indicating elevation (A-G for 40° to −20° degrees elevation) and numbers indicating azimuth (1-19 indicating 90° to −90° azimuth). Participant responses were collected on an Apple iPad held by the participant during the experiment. Participants were instructed to direct their head at the loudspeaker directly in front of them for the duration of the stimulus. Once the stimulus ended, participants could look at the iPad to enter their response. Participants were instructed to redirect their head towards the front loudspeaker before triggering the start of the next trial. Compliance with these instructions was confirmed by experimenter observation.

#### Procedure

The trial procedure and task were identical to that of Experiment 1 (which was run online), except that participants heard sounds from loudspeakers rather than headphones/earphones, and that the cue/target/distractor varied in spatial position. First, a 2-second cue excerpt would play from a specific location, allowing participants to hear both the target talker’s voice and location. After a 0.5-second delay, a 2-second target excerpt was played from the same location, while two 2-second distractor excerpts were played simultaneously at locations that varied with respect to the target location. The target and distractor were separated in either azimuth or elevation, depending on the condition.

Two distractors were used in every trial. When the distractors varied in azimuth, the two (symmetrically placed) distractors served to preserve the signal-to-noise ratio at each ear across different target-distractor separations^34,113^. When the distractors varied in elevation, the use of two distractors served to match the distractor properties to those of the azimuth conditions, enabling a more controlled comparison. Listeners reported the middle word spoken by the target talker (i.e. the word overlapping the 1-second mark). Participants entered responses by typing on an iPad, connected to a local server running the same experimental interface used in the online experiments.

A common set of target positions and distractor offsets were used in azimuth and elevation conditions to compare the effect of spatial separation in each dimension. Distractors were offset relative to target positions by either 0 (i.e., co-located), 10, or 60 degrees in either azimuth or elevation. These offsets were chosen as those likely to reveal the extent of spatial tuning subject to the limits of our speaker array: 10 degrees was the smallest offset between speakers, while 60 degrees was the largest span available in elevation (minimum elevation of −20 degrees; maximum of 40 degrees). Cue and target excerpts were always presented at 0 azimuth, and at either −20- or 40-degrees elevation (so that 60-degree target-distractor offsets could be used). To equate the uncertainty of the target location in azimuth and elevation, one of the two possible target elevations was sampled randomly for each participant, and was used for all trials (so that for any single participant the target was always at the same location in azimuth and elevation).

In the conditions in which target and distractors were separated in azimuth, the cue, target, and distractor signals shared a common elevation. The two distractor excerpts were presented symmetrically around the target at the specified offset. In the conditions in which target and distractors were separated in elevation, cue and target excerpts were always presented at 0 degrees azimuth. The two distractor excerpts were presented from the same position, at 0 azimuth and an elevation offset from the target elevation by the specified amount. When the target was at −20 degrees the distractors were offset above the target location; when the target was at 40 degrees the distractors were offset below the target location.

The experiment used the method of constant stimuli. At each target-distractor position, word recognition was measured at a set of signal-to-noise ratios (−9, −6, −3, 0, 3, and 6 dB). In trials with co-located distractors, the two distractor waveforms were first summed, and then their combined signal was normalized to 65 dB SPL. In trials with symmetrically positioned distractors, each distractor was individually set to 61.99 dB SPL, such that the level of the two distractors was 65 dB SPL on average when summed at the ear. The level of the target excerpt was then adjusted to obtain the desired SNR as 65 dB + SNR. The cue signal was always presented at 65 dB SPL, so that cue level would not be informative for the task.

Participants completed 16 trials for each combination of target position, distractor offset, and SNR, totaling 480 trials (1 target position x 5 distractor offsets x 6 SNRs x 16 trials). Trials were randomly ordered and then grouped into six blocks of 80 trials, with participants encouraged to take self-timed breaks after each block. The entire experiment took approximately 2-hours to complete. The selection of stimuli and assignment of excerpt to condition were randomized independently for each participant. 33 participants (22 female, 10 male) completed the experiment. Participant ages were between 18 and 40 years (median age = 23).

#### Analysis

Speech reception thresholds were estimated for each distractor offset in azimuth or elevation using the following procedure. First, word recognition performance as a function of signal-to-noise ratio was measured for each participant as in Experiment 1, at each distractor offset. Second, thresholds at each distractor offset and offset direction were estimated by fitting a second-order polynomial to the mean of these values across participants and calculating the SNR granting 50% performance from this curve. The uncertainty in these thresholds was estimated via bootstrap (thresholds were measured from 10,000 bootstrap samples of 33 participants; results graphs plot the mean and standard deviation of these thresholds).

#### Model experiment

Models were tested at all combinations of the 1,952 stimuli (all cue, target, and two-distractor pairings) with the 5 distractor offsets and 6 SNRs (58,560 total combinations). Stimuli were spatialized using a virtual rendering of the loudspeaker array room participants were tested in. Sound pressure levels in the model experiment were identical to the human experiment. To approximately equate the model and human testing conditions, 30dB of pink noise was added to the model stimuli to match the background noise level present in the loudspeaker array room used to test participants. Model speech reception thresholds for each distractor offset and direction were estimated by bootstrapping over the 10 model architectures (10,000 samples).

### Experiment 7: measuring the width of spatial attention

Experiment 7 measured how spatial release from masking in azimuth depends on the azimuthal position of the attended (target) talker relative to the listener. Eight spatial configurations were used: 2 target azimuths (0° or 90°) x 4 distractor azimuthal offsets (0°, 10°, 30°, or 90°). Experiment 7 used the same cue, target, and distractor clips as Experiment 6, except that only one distractor talker was included on each trial. All stimuli were presented at 65 dB SPL, with the target and distractor signal-to-noise ratio set to 0 dB. Participant and model responses were analyzed as in Experiment 1.

#### Procedure

Experiment 7 used the same procedure as Experiment 6, except for the following changes. Participants completed 160 total trials (2 target positions x 4 distractor offsets x 20 trials). Each trial had a different target word. The 160 target words for a participant were randomly sampled from the full set of 488 target words, and their assignment to spatial locations was randomized per participant. Trials were arranged into 4 blocks of 40 trials. All the trials in a block had the same target position, and the block included 10 trials of each distractor offset. The block order and the order of trials within blocks was randomized per participant. 28 participants (16 female, 12 male) completed the experiment. Participant ages were between 19 and 39 years (median age = 26).

Because the speaker array did not enable left-right symmetric offsets at the 90° azimuth positions (the array did not extend beyond 90° on either side), we used the following procedure to better equate the uncertainty over source positions between the 0° and 90° target conditions. First, distractors were offset in a consistent direction relative to the target (i.e., distractors occurred either to the left or right of the target for the entire experiment). The distractor offset direction was alternated across participants. Second, the 0° and 90° target conditions were achieved by rotating participants to face either the center or the side of the array, while maintaining the source playback locations. In all trials, the cue and target were presented at the center of the array, with distractors occurring 0°, 10°, 30°, or 90° away from the target in azimuth (either to the participant’s left or right depending on the offset direction). In blocks testing the 0° target condition, participants faced the center speaker. In blocks testing the 90° target condition, participants were rotated so the center speaker was at 90° relative to the participant, and the distractor offsets were positioned to the same side as in the 0° target condition. All stimuli were presented at 0° elevation.

#### Model experiment

Models were tested at all combinations of the 1,952 stimuli (cue, target, and single-distractor pairs) with the 2 target positions and 4 distractor offsets (15,616 total combinations). All model stimuli were RMS normalized to 0.02. The room simulation and inclusion of noise were identical to Experiment 6.

### Analysis of model locus of attention (Figure 5)

To analyze the locus of attention in our models, we compared the model’s representation of a single-talker target clip with its representation of a mixture composed of the target and a single-talker distractor clip, as a function of whether the model was cued to the target talker or distractor talker. The logic for this analysis was that the representations of a single source and a mixture containing that source would be similar if attentional selection for the single source’s talker was applied to the mixture. We analyzed the similarity between targets and mixtures containing one distractor talker, presented signals diotically to the model, and quantified similarity using Pearson’s correlation coefficient. The cue, target, and single-distractor speech excerpts curated for Experiment 1 were used as stimuli. To measure any effect of spatial separation on this analysis, we included three azimuthal separation conditions: 0°, 10° (e.g. target at +5° and distractor at −5° or vice versa), and 90° (e.g. target at +45° and distractor at −45° or vice versa) separation. Activations for all excerpt pairs in each azimuthal separation condition were measured. The following procedure was performed for each cue, target, and one-distractor pair in each separation condition. First, representations of the isolated target clip, 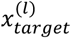, distractor clip, 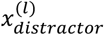, and their mixture, 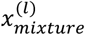, were obtained separately from each layer, *l*, using the same cue to designate the target talker for each. Second, we measured Pearson’s correlation coefficient between target and mixture as corr : 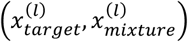, and between distractor and mixture as corr : 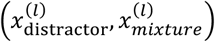 where corr(*x, y*) is the Pearson’s correlation:

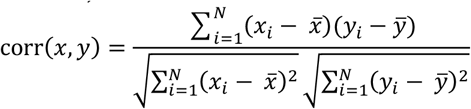

and *i* indexes the flattened feature dimension of size *N* = Channels × Frequency × Time. 1000 cue, target, and single-distractor pairs were used to measure correlations (500 same-sex and 500 different-sex distractors). Targets and distractors were presented at 0 dB SNR, and all signals were RMS-normalized to 0.02.

### Analysis of model feature tuning (Supplementary Figure 11)

To analyze feature tuning, we measured model unit activations to a large set of spatialized speech excerpts from the best performing feature-gain model. We used 5,000 speech excerpts from the Word-Speaker-Noise dataset^94,99^ as this dataset had a similar vocabulary to our training task, but was constructed so as to balance talkers across word classes. We assessed whether unit responses could be explained by the pitch, location, word, or talker of a speech excerpt. We used the average F0 of the voiced segments of an excerpt (estimated using pyin^114^) to capture pitch. Pitch was discretized into 54 bins, determined by calculating the appropriate bin width using the Freedman-Diaconis method:

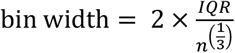

where IQR is the interquartile range over average F0 and *n* is the number of excerpts. We spatialized speech excerpts to 0°, ±10°, ±45°, and 90° azimuth and 0° elevation for 7 total locations. The subset of the Word-Speaker-Noise dataset that we used contained 431 talkers and 776 words. All of the 5,000 speech excerpts were spatialized to each of the 7 locations. We then ran four one-factor ANOVAs on each unit’s responses, one for each of the four variables, and calculated the variance explained by each variable from the sums of squares. This analysis was performed using the ordinary least squares and linear model ANOVA methods of the statsmodels^115^ python package. We report the mean variance explained for each feature, at each model stage.

### Aggregate measures of human-model similarity (Figure 6b&c)

Human-model behavioral similarity was quantified by comparing human and model performance in each experimental condition, separately for each model. To ensure that our conclusions were robust to the choice of similarity metric, the following analysis was performed using Pearson’s correlation coefficient root-mean-squared-error (RMSE), and mutual information as similarity metrics. For each model, we measured the similarity between the mean human behavior (averaged across experiment participants) and mean model behavior, using means computed per experimental condition (e.g. a particular distractor type x signal-to-noise ratio). For the feature-gain model, the mean model behavior was taken as the average across the 10 network architectures. To calculate mutual information, we generated histograms of human and model performance. To ensure robustness to histogram bin size, mutual information was computed using bin counts of 5, 10, 15, and 20 bins. Error bars on the mutual information for each bin size were obtained via bootstrap (1000 samples).

The statistical significance of the effect each alternative architectural constraint (e.g. early-only, late-only, or attention-free architectures) had on overall human-model similarity was assessed by comparing the human-model similarity scores against the distribution of the feature-gain model’s human-model similarity scores via sign test. Two-tailed p values are reported, and effect sizes were quantified by measuring the difference between similarity scores over the mean score of the feature-gain model.

### Representational dissimilarity analysis of spatial cues (Supplementary figure 7)

We quantified acoustic spatial cues by calculating the difference between the binaural cochlear representations of the same speech excerpt presented at different spatial positions. Specifically, we generated representational dissimilarity matrices (RDMs) for a) pairs of single target speech excerpts rendered at different positions, as a function of their spatial separation in either elevation or azimuth, and b) pairs of single target speech excerpts and the mixture of that target speech excerpt with a distractor excerpt. In the second case the distractor was always presented at the midline. The rationale was to test 1) whether the presence of the distractor disrupted spatial cues of the target and 2) whether there was evidence that spatial cues were less precise in the periphery compared to the midline. Dissimilarity was computed as 1 − *r*_*xy*_, where *r*_*xy*_ is Pearson’s correlation between the binaural cochleagrams of signals *x* and *y*. In each RDM, the *ij*_*th*_ entry was computed as the average dissimilarity between representations of signals whose sources were rendered at location *i* and location *j*. To measure dissimilarity in azimuth, target sources were spatially rendered from 90° to −90° azimuth (in 10° increments) and 0° elevation. To measure dissimilarity in elevation, target sources were spatially rendered from −20° to 40° elevation (in 10° increments) and 0° azimuth. Speech excerpts were drawn from the set of 500 target and same-sex distractors used in the stage of selection analysis, originally cut for Experiment 1.

### Statistics

Statistical tests: Interactions were assessed using a repeated measures Analysis of variance (ANOVA), implemented using the statsmodels^115^ and Pingouin^116^ packages for python. Significance of the effect of model architectural constrains on human-model similarity used sign tests implemented via statsmodels and scipy.stats packages. Paired two-tailed t-tests used to measure the effect of distractor harmonicity on model performance were implemented using the scipy.stats^117^ packages.

Experiment 6 did not permit a conventional test for an interaction because thresholds were difficult to estimate reliably in individual participants. Instead, we assessed the statistical significance of the interaction between the direction of offset (azimuth or elevation) and offset magnitude (0°, 10°, or 60°) using a permutation test. First, the interaction between conditions was measured by subtracting off the marginal contribution of each independent variable:

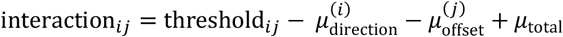

where *i* indexes the direction of offset (azimuth or elevation), *j* indexes the offset magnitude, 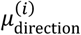 is the average threshold over offsets for direction *i*, 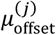 is the average threshold over directions for offset *j*, and *μ*_total_ is the average over both direction and offset. An overall interaction effect was then obtained as the sum of squares of these interaction terms:

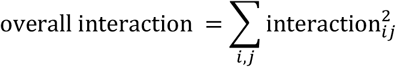

The overall interaction was re-computed 10,000 times with direction of offset and offset magnitude labels permuted at the subject level, obtaining a null distribution used to calculate a *P*-value for the actual overall interaction.

Error bars: Except where otherwise noted in figure captions, error bars in results figures indicate ±1 standard error of the mean (SEM) across experiment participant results (human results) or across the 10 network architectures (model results).

## Data availability

The data that support the findings of this study are available at the project repository.

## Code availability

The code associated with this study is available at the project repository.

## Acknowledgments

Research supported by National Institutes of Health grant R01 DC017970. The funders had no role in study design, data collection and analysis, decision to publish or preparation of the manuscript. The authors thank Adanna Thomas for running the human participants in Experiments 6 and 7.

## Author contributions

I.G. and J.H.M. conceived the project and wrote the first draft of the paper. I.G. and R.P.H. trained the models, designed the experiments, and ran the experiments. I.G. made the figures. All authors edited the paper.

## Competing interests

The authors declare no competing interests.

## Supplementary Figures and Tables for Griffith, Hess & McDermott

**Supplementary Fig. 1.**
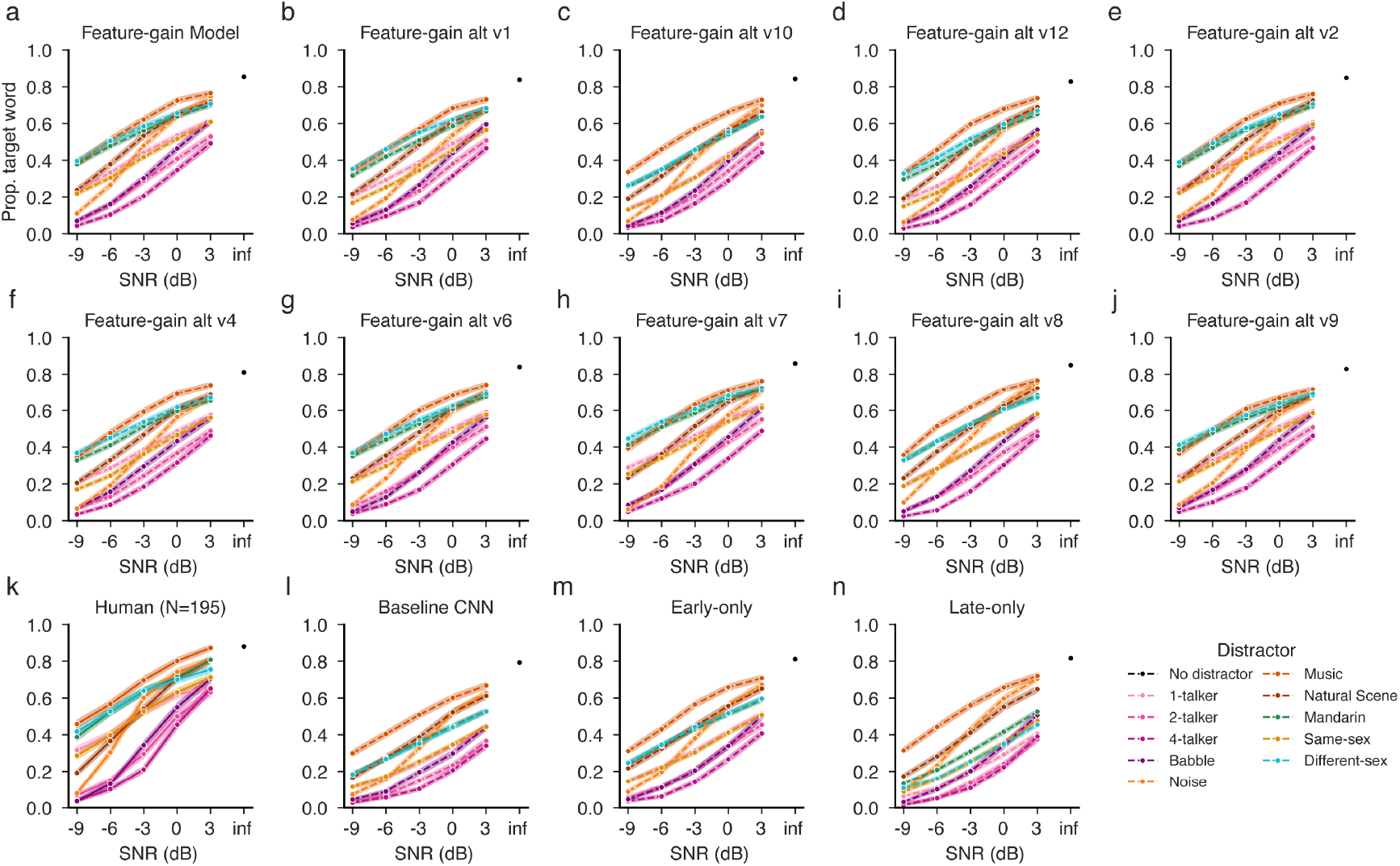
Performance of all models on Experiment 1. Word recognition as a function of signal-to-noise ratio and distractor type for each model architecture. Panels a-j show results for each individual feature-gain model architecture (main figures show results averaged across these ten architectures, with different conditions plotted separately to elucidate particular effects of interest). Panel k shows results for human participants, replotted from Figure 2. Panels l-n show the baseline, early-only, and late-only architectures which each perform worse with speech-on-speech examples than both humans and the feature-gain models, particularly with 1-distractor (light-pink, gold, green, and blue lines).

**Supplementary Fig. 2.**
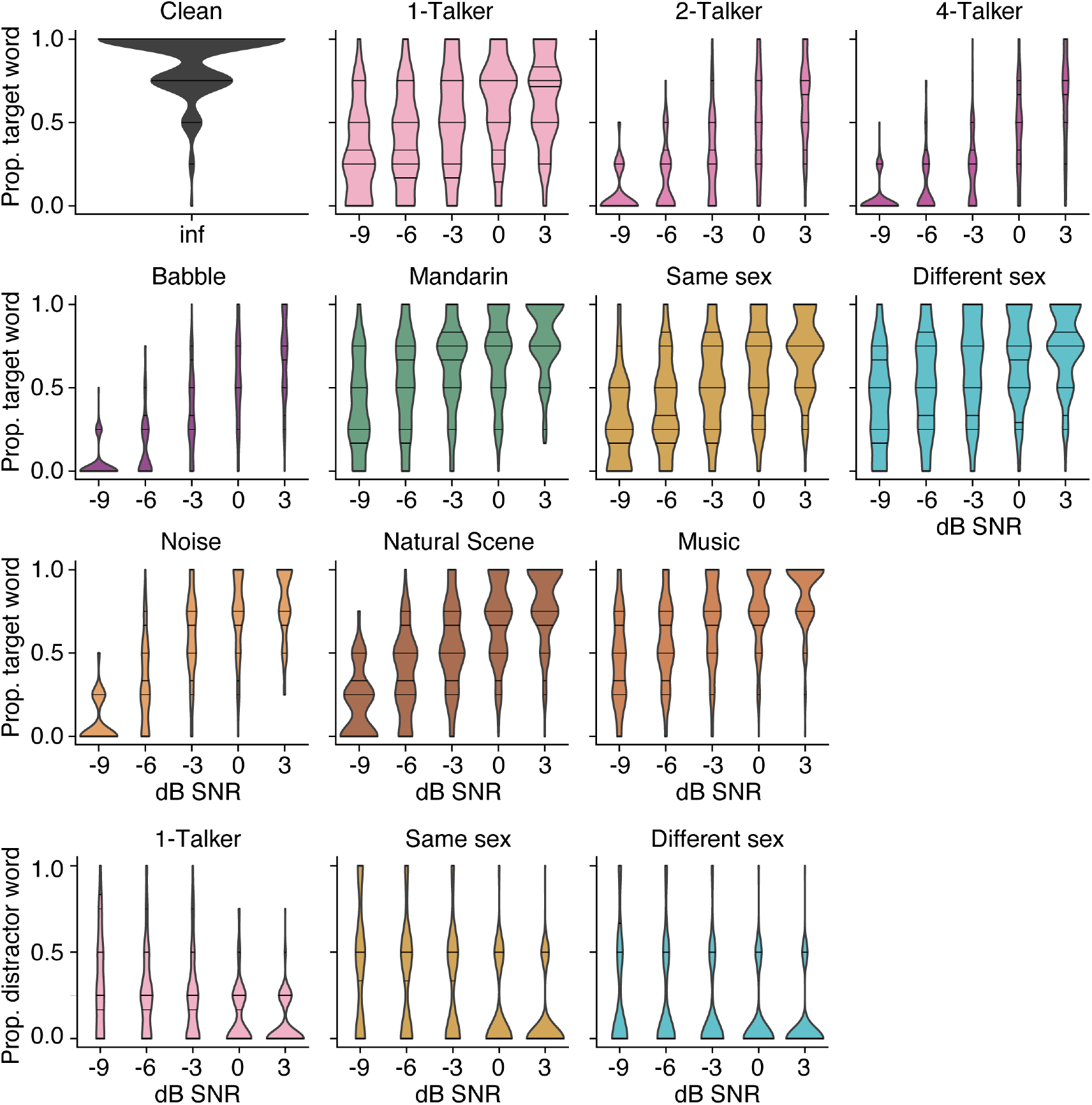
Data distributions for Experiment 1. Results of Experiment 1 plotted as distributions of the performance of individual participants. This figure plots the same results as Figure 2a-e, but separates each stimulus condition into a separate panel so that the data distributions can be shown.

**Supplementary Fig. 3.**
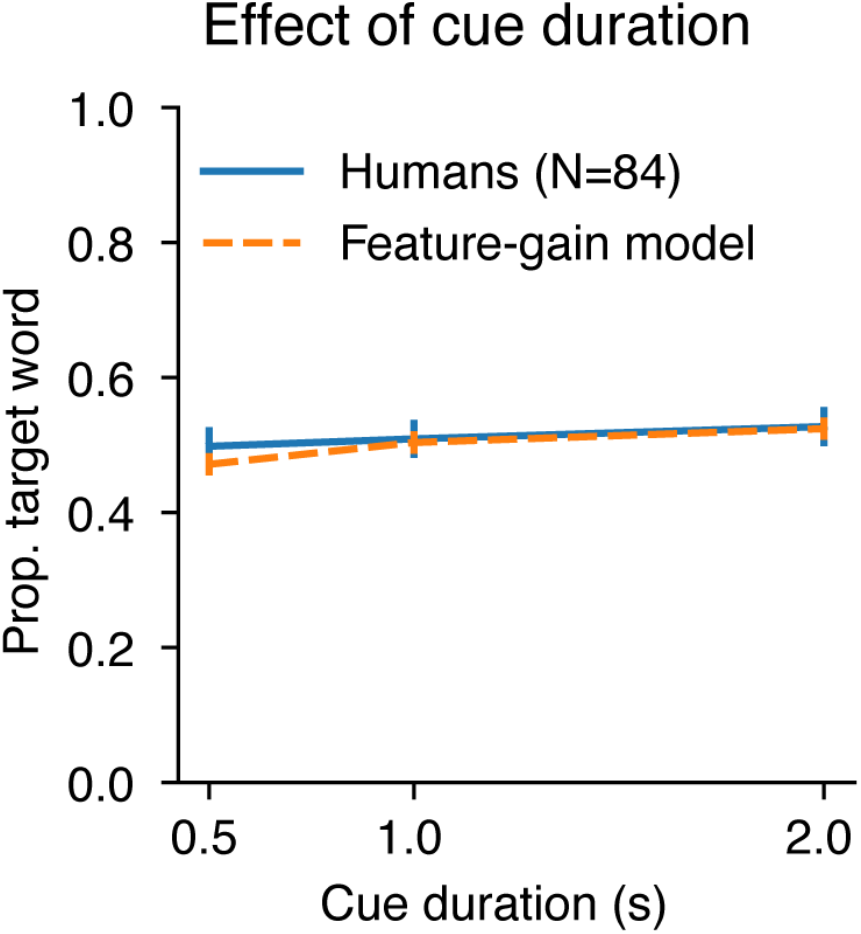
Human and model performance for different cue durations. Results for Experiment 1b, which measured recognition of words spoken by a cued target voice superimposed on a single distractor talker at 0 dB SNR. Error bars plot standard error.

**Supplementary Fig. 4.**
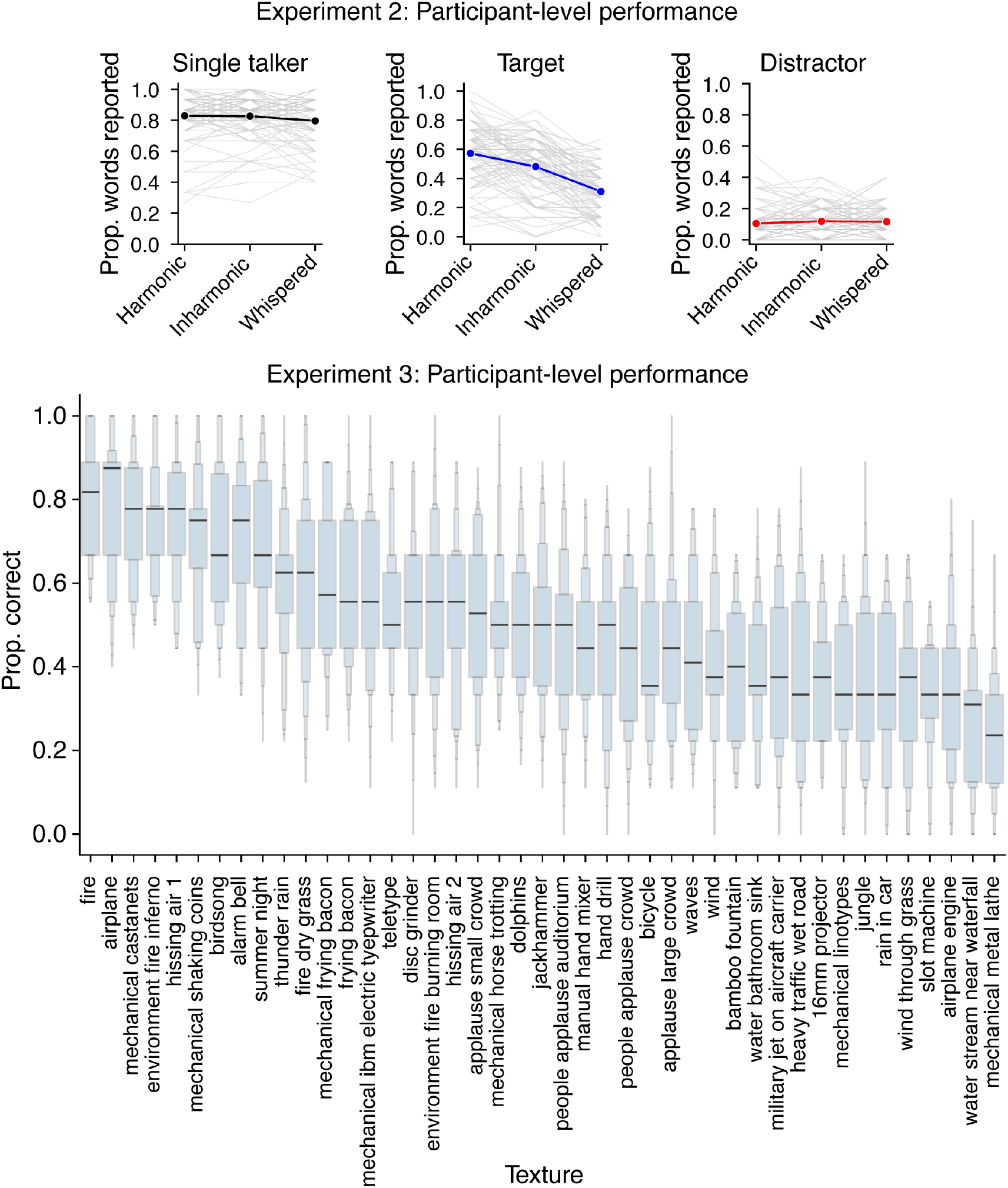
Data distributions for Experiments 2 and 3. Results of Experiments 2 and 3 plotted as distributions of the performance of individual participants. This figure plots the same results as Figure 2f&h, but separates each stimulus condition into a separate panel so that the data distributions can be shown.

**Supplementary Fig. 5.**
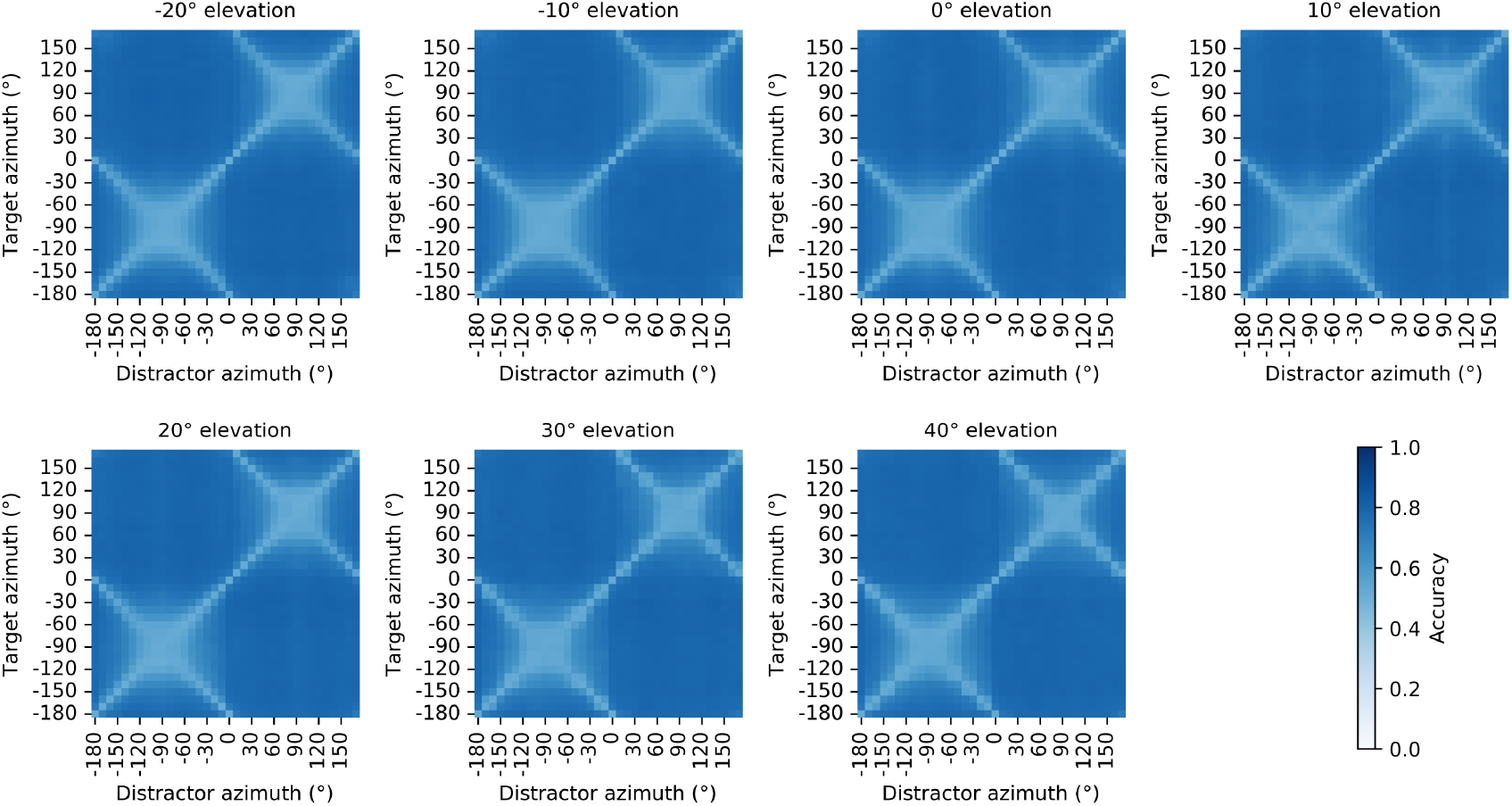
Model word recognition at all target-distractor azimuth pairings. Word recognition performance for the main feature-gain model (arch_v00 in Supplementary Table 1) as a function of target and distractor positions in azimuth. Each panel plots results for a different elevation, with target azimuth on the vertical axis and distractor azimuth on the horizontal axis (targets and distractors were at the same elevation). The plots differ from those in Figure 4 in plotting results for the full 360 degrees of azimuthal positions (for completeness). Because there was little effect of whether a source was in the front or back hemisphere, the plots in Figure 4 averaged the two hemispheres. The front-back symmetry yields the “X” structures evident in the plots. The broadening of spatial acuity as the target azimuth moved peripherally (i.e. approaching ±90° from 0°) was apparent at all elevations.

**Supplementary Fig. 6.**
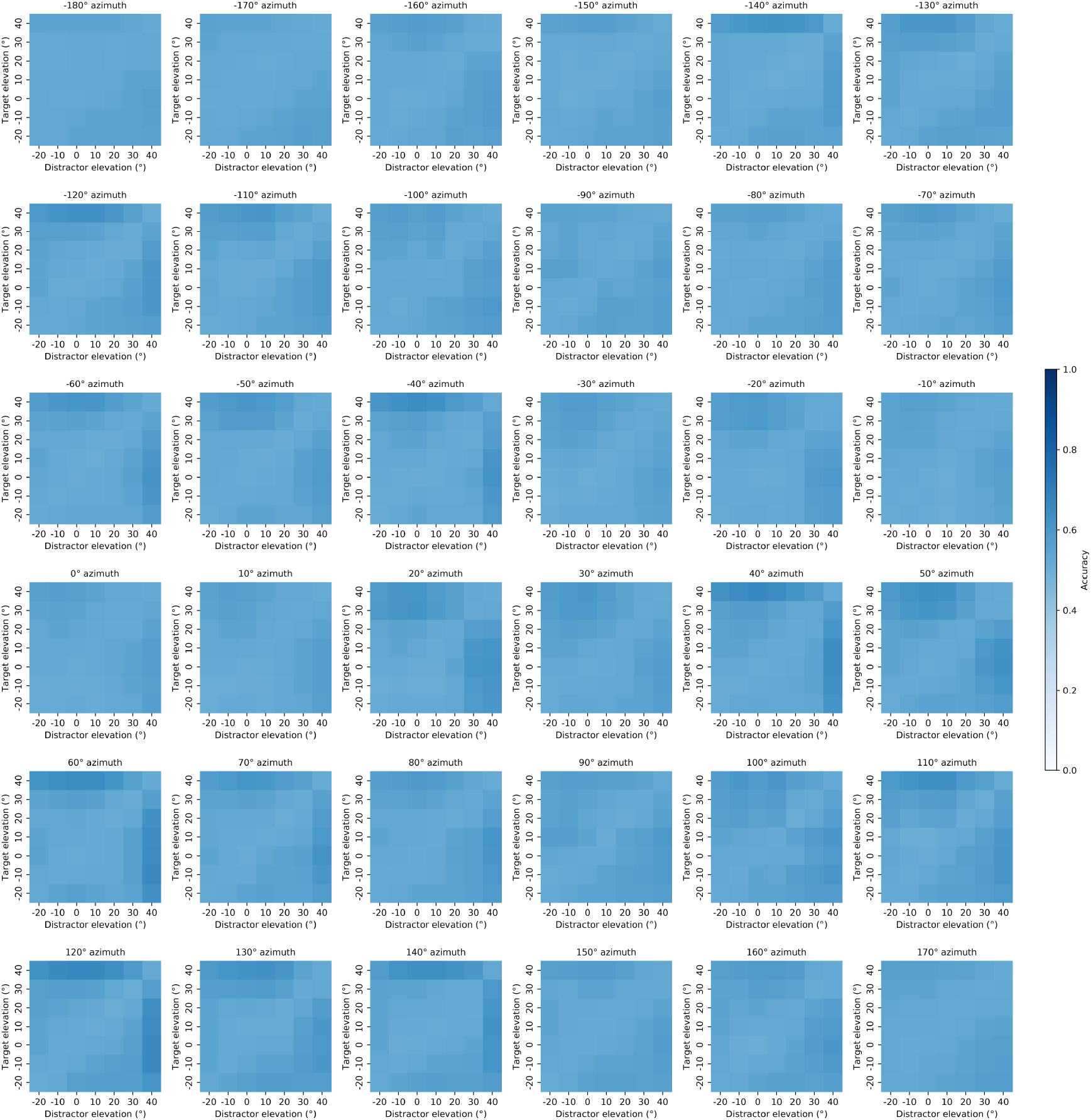
Model word recognition at all target-distractor elevation pairings. Word recognition performance for the main feature gain model (arch_v00 in Supplementary Table 1) as a function of target and distractor positions in elevation. Each panel plots results for a different azimuth, with target elevation on the vertical axis and distractor elevation on the horizontal axis (targets and distractors were at the same azimuth). Target-distractor offset in elevation produced little benefit regardless of azimuthal position.

**Supplementary Fig. 7.**
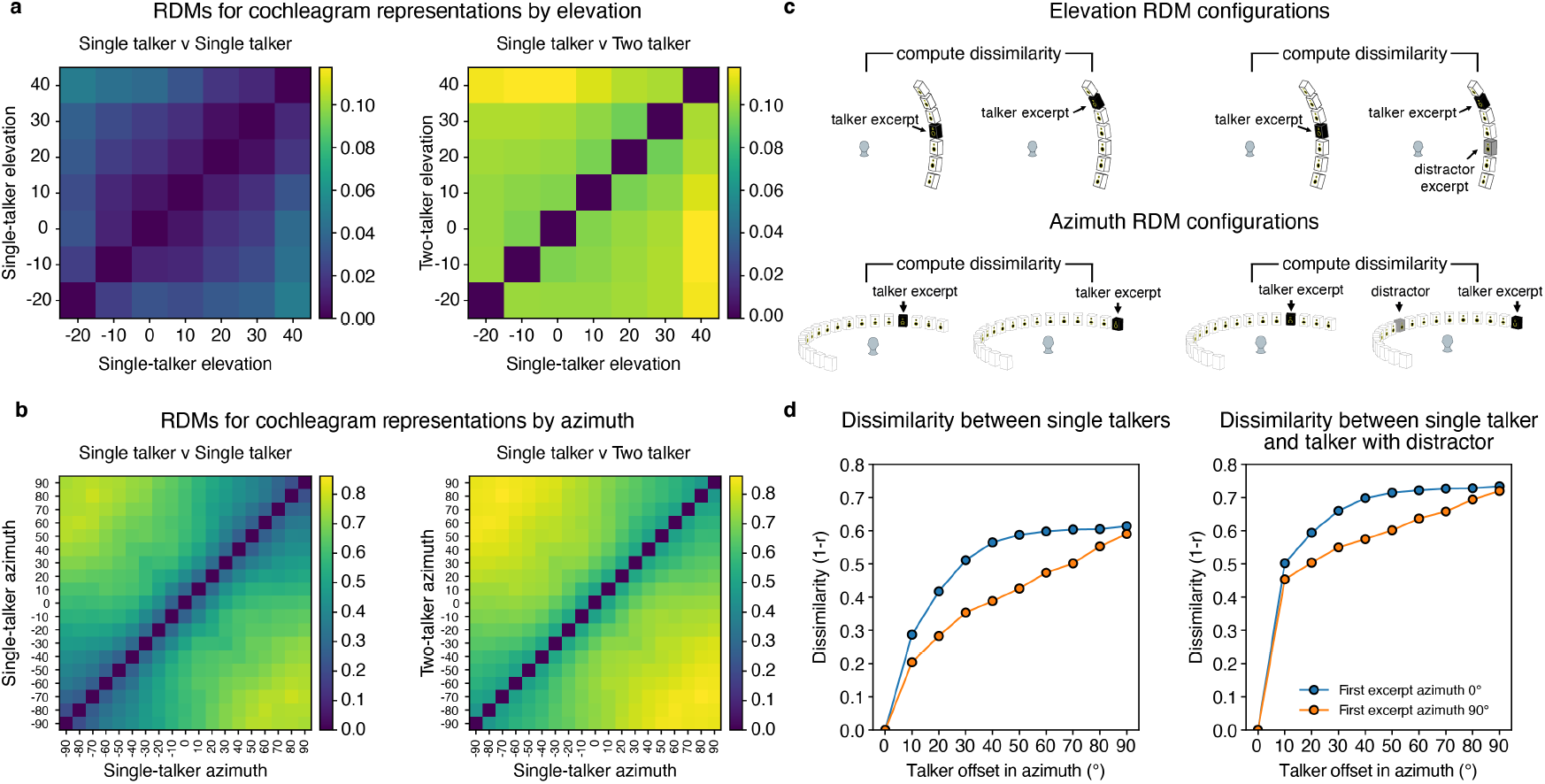
Effect of concurrent talker on spatial cues. We quantified spatial cues to azimuth and elevation by calculating the dissimilarity between the cochleagram representations of different stimuli as a function of differences in the spatial positions of speech signals rendered in the stimuli. We first measured this dissimilarity for the same speech signal rendered at different positions. We used the same speech signal so that interaural time differences would be reflected in the cochleagram correlation (if the speech signals had not been identical, their fine structure would have been uncorrelated irrespective of any spatial cue from the interaural time difference). We then measured the dissimilarity with a distractor talker superimposed on the second signal. The distractor was positioned in front of the listener (at zero elevation). We expected to see a gradient of dissimilarity as the spatial offset was increased, and that this might be reduced by the distractor talker. **a**. Results for elevation offset. A gradient of dissimilarity is evident as talker offset is increased. This gradient is less evidence in the presence of a distractor talker. Dissimilarity was measured using the time-averaged cochleagram, to best reveal spectral cues to elevation. **b**. Results for azimuth offset. A gradient of dissimilarity is evident even in the presence of a distractor talker, indicating that azimuth cues are robust to the presence of a concurrent talker. **c**. Schematic of the elevation (top row) and azimuth (bottom row) configurations included in the dissimilarity analysis. Dissimilarity was computed either between representations of single talkers (left column), or between representations of a single talker and representations of a talker with a concurrent distractor (right column). **d**. Comparison of dissimilarity gradient for a target talker at the midline (0 degrees) vs. at the periphery (90 degrees). The dissimilarity increases rapidly for the talker at the midline, and more gradually for the talker at the periphery, consistent with the idea that binaural spatial cues permit a tighter spatial attentional focus at the midline compared to the periphery.

**Supplementary Fig. 8.**
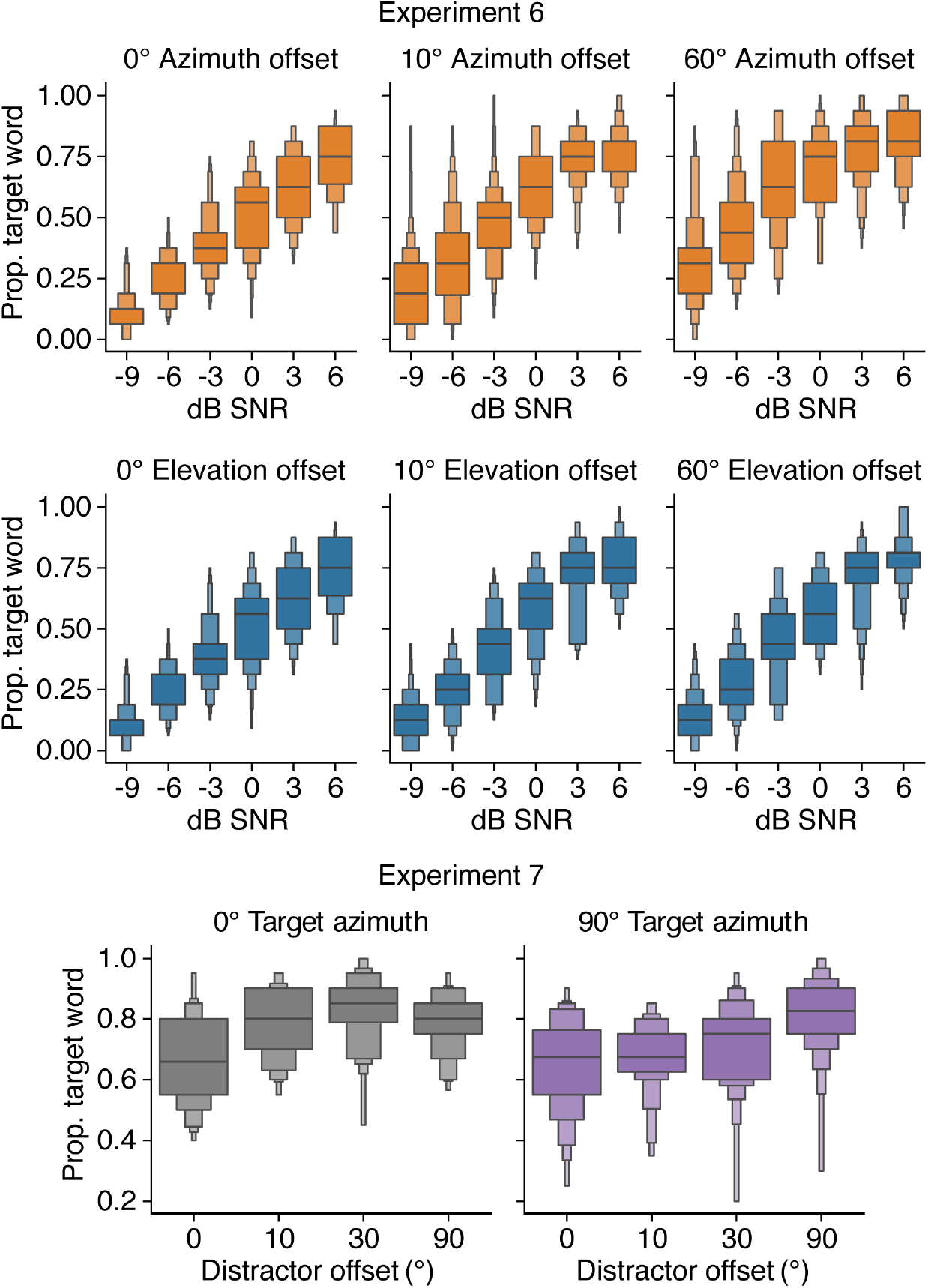
Data distributions for Experiments 6 and 7. Results of Experiments 6 and 7 plotted as distributions of the performance of individual participants. This figure plots the same results as Figure 4c&e, but separates each stimulus condition into a separate panel so that the data distributions can be shown.

**Supplementary Fig. 9.**
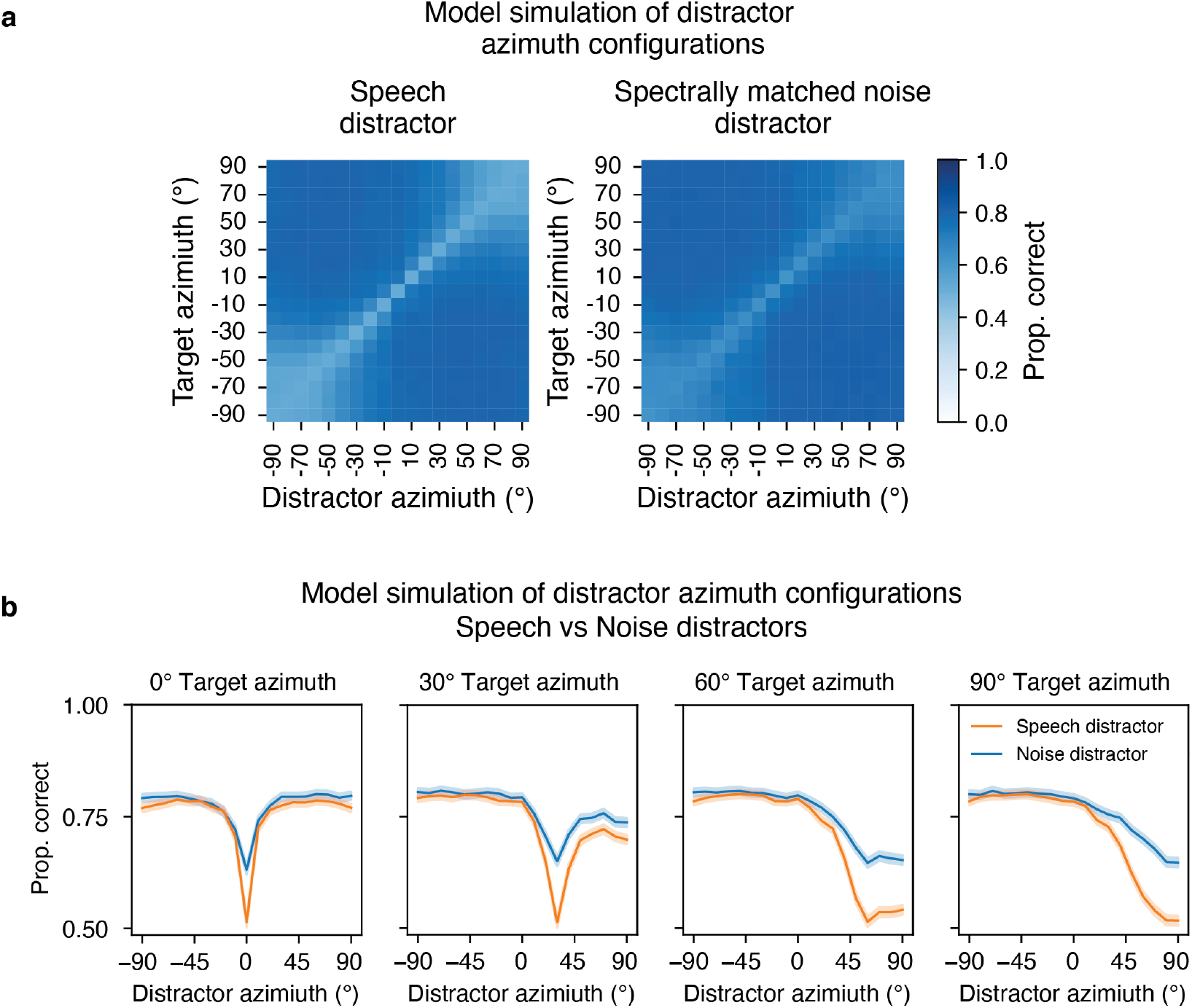
Effect of target and distractor spatial position on model performance for speech and noise distractors. **a**. Word recognition performance for the main feature-gain model (arch_v00 in Supplementary Table 1) is plotted as a function of target and distractor positions in azimuth, for both single-talker distractors and speech-shaped noise distractors. As in Figure 4, results are averaged across front and back positions for ease of visualization. **b**. Data from (a) plotted as line graphs for four example target positions, to enable direct comparison of performance for speech and noise distractors. Shaded regions plot confidence intervals (95%) obtained via bootstrap. Performance with noise distractors varies much less with spatial position than performance with speech distractors, as in humans^118,119^. The presumptive explanation is that voice features alone are sufficient to select speech from noise, such that there is less benefit from spatial attention. There is nonetheless some spatial benefit on performance, consistent with prior results in human listeners^119^. This spatial dependence could be explained by a combination of changes in signal-to-noise ratio at the ear (which improves at the ear closer to the target as the distractor is moved away) and binaural effects related to masking^120,121^. However, the larger effect for speech distractors suggests a contribution of spatial attention in this setting. The similar spatial dependence for speech and noise distractors presumably reflects the spatial dependence of the underlying binaural cues, which also cause localization acuity to be worse in the periphery.

**Supplementary Fig. 10.**
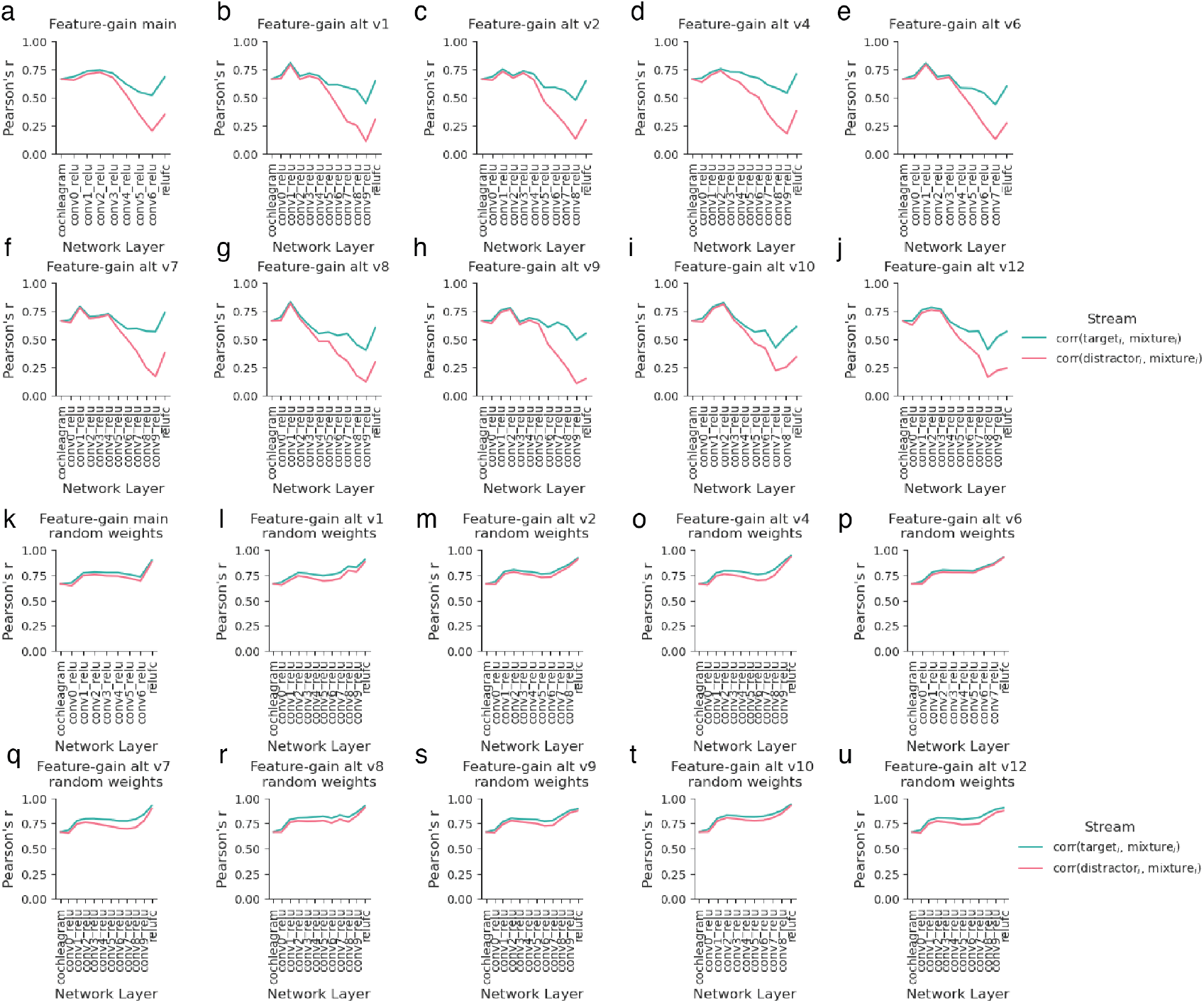
Stage-of-selection analysis for each model architecture. Analysis of Fig. 5, plotted separately for each model architecture. All feature-gain architectures displayed fairly similar patterns of target enhancement at late model stages (panels a-j). Each architecture initialized with random weights did not show the same trend (panels k-u).

**Supplementary Fig. 11.**
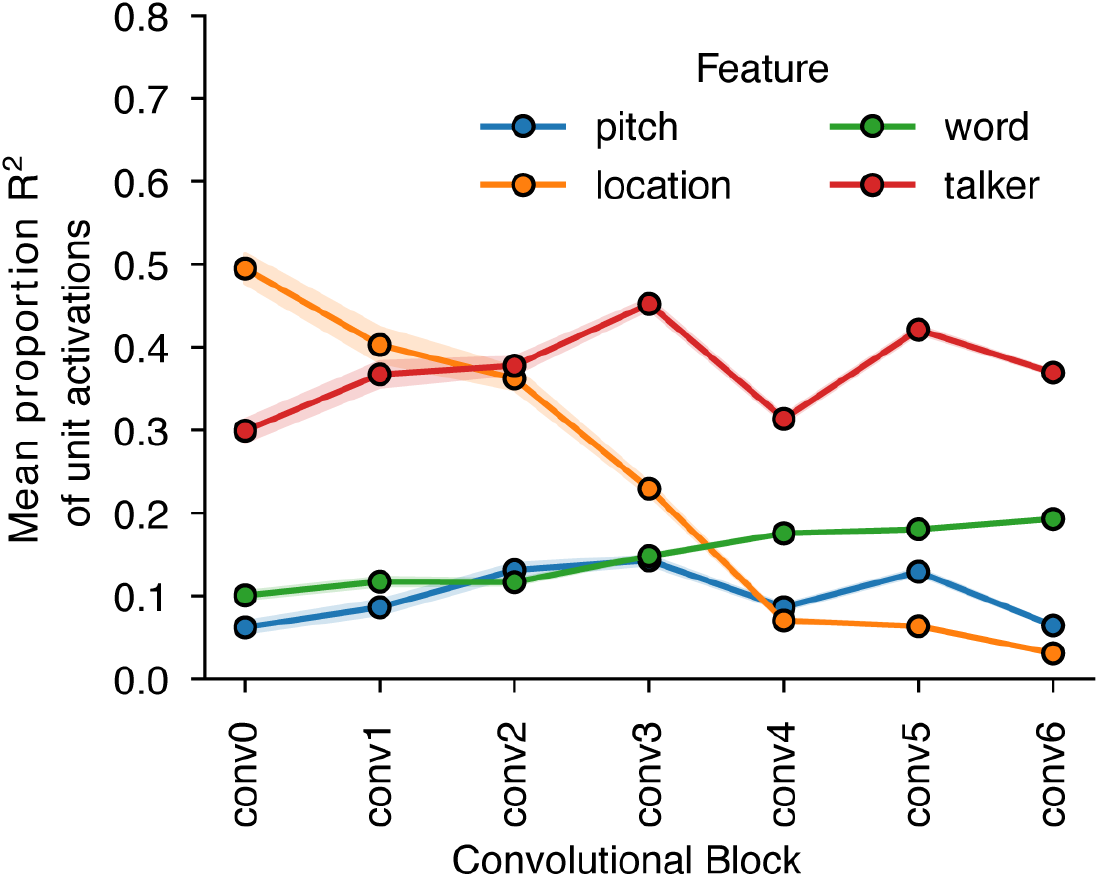
Feature tuning across model stages. We measured model activations for a set of speech excerpts rendered at different spatial locations, and analyzed the variance of each unit’s response that was explained by location, pitch, talker, and the middle word of the excerpt. The graph plots the mean of this quantity at each model stage. The two dimensions that are useful for selection (location and talker) account for considerable response variance in early model stages. Tuning to words is weak in early stages and increases towards the end of the model. Tuning to pitch (specifically, the mean f0 of the voiced segments of the speech excerpt), which is one of many features defining the talker’s voice, is evident throughout the model. Location tuning was less prevalent in deep model stages, whereas talker tuning persisted, potentially because talker cues are extracted over a larger number of hierarchical stages.

**Supplementary Fig. 12.**
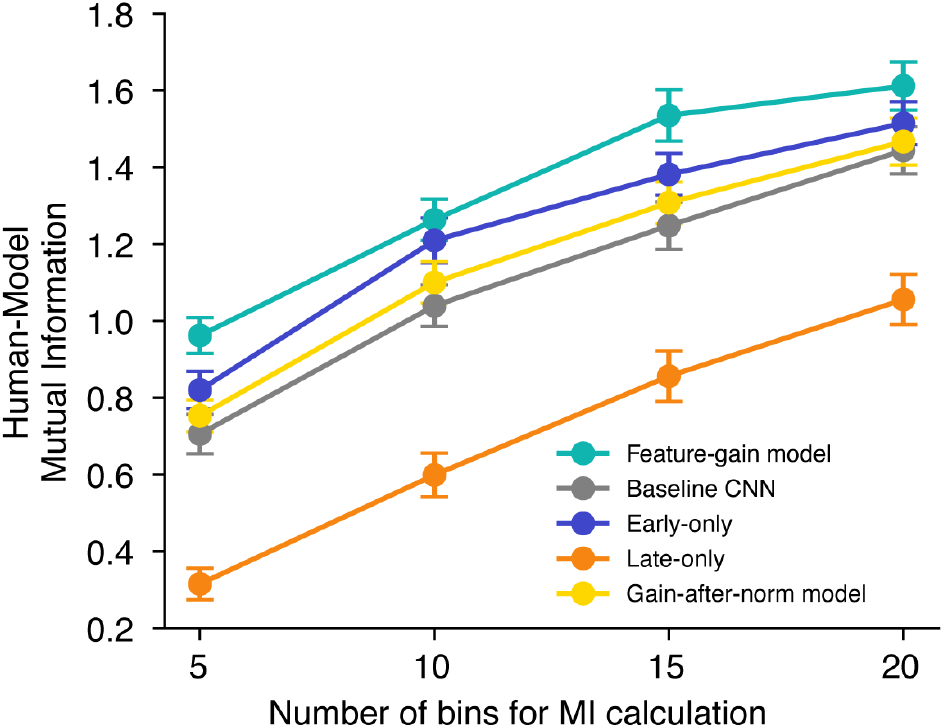
Analysis of human-model similarity using mutual information. Mutual information was calculated using histograms of human and model proportion correct in different experimental conditions. We performed the analysis for different numbers of histogram bins to ensure that the results were robust to the bin width. Error bars plot standard error, derived via bootstrap.

**Supplementary Fig. 13.**
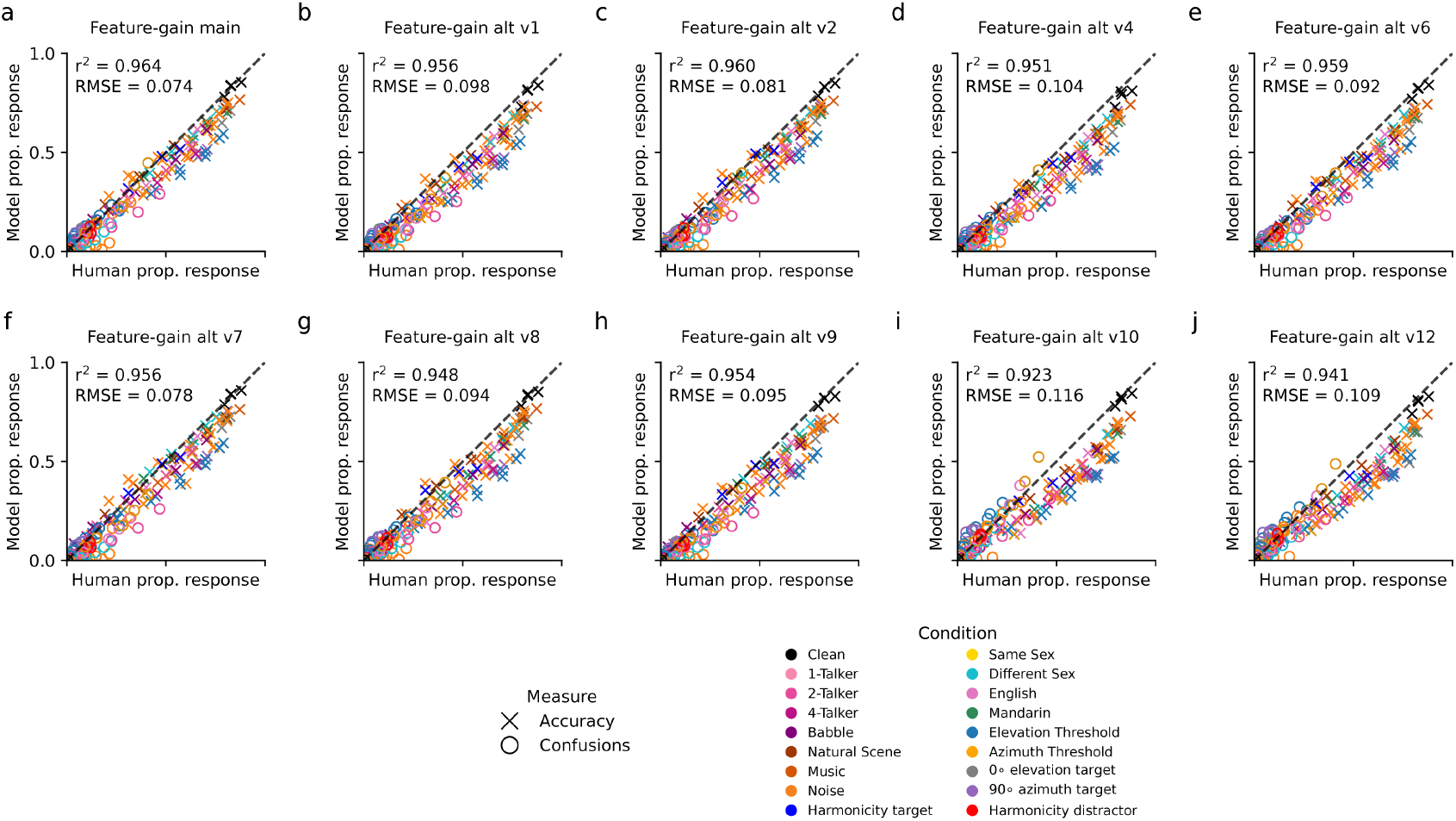
Comparison of human and model performance for each model architecture. Scatter plots of model vs. human performance for each individual feature-gain architecture.

**Supplementary Fig. 14.**
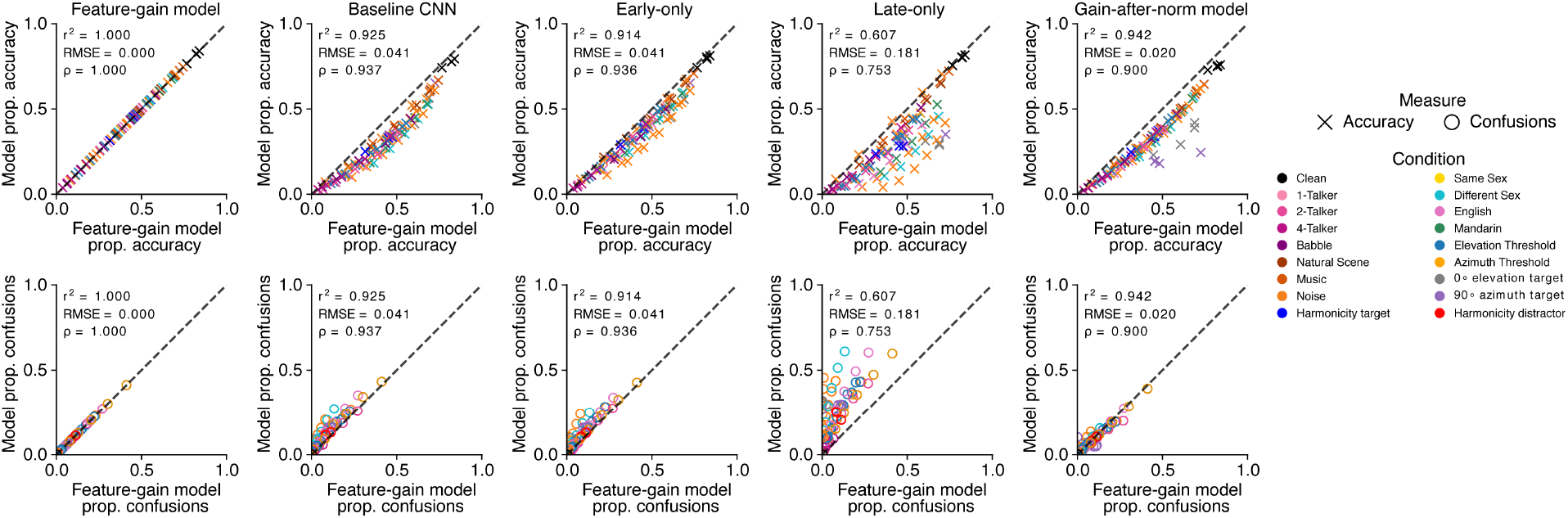
Comparison of performance of different model architectures. Scatter plots of alternative model performance vs. that of the feature-gain model.

**Supplementary Table 1.**
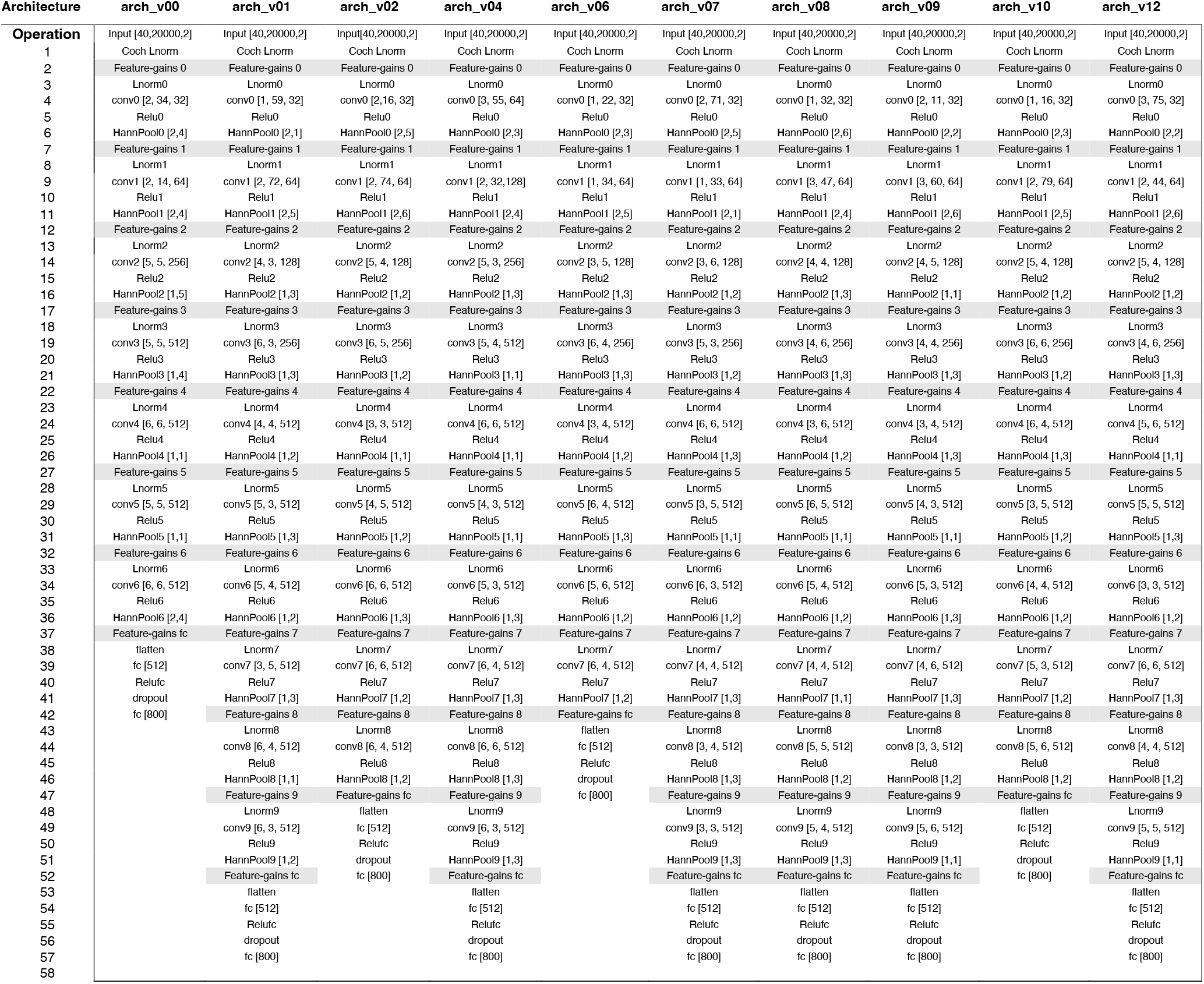
Neural network architectures for feature-gain models. Grey bands indicate stages where feature-gain operations occurred. Legend:

- *Lnorm*: layer normalization operation
- conv [*h, w, k*] : convolutional layer with *h* = kernel height (frequency dimension), *w* = kernel width (time dimension), and *k* = number of kernels
- Relu: rectified linear unit activation function
- HannPool [*s*_*f*_, *s*_*t*_] : Hanning window weighted averaging pooling operation with stride *s*_*f*_in the frequency dimension and stride *s*_*t*_ in the time dimension
- flatten: reshape operation to map a multidimensional tensor to a vector
- dropout: dropout regularization with 50% dropout rate
- fc [*N*]: fully-connected layer with *N* units

## Notes

### Competing Interest Statement

The authors have declared no competing interest.

### Summary of Updates

Figures revised; Supplemental figures updated; text revised clarify corresponding figure changes.

